# Fast Adaptation to Rule Switching using Neuronal Surprise

**DOI:** 10.1101/2022.09.13.507727

**Authors:** Martin Barry, Wulfram Gerstner

## Abstract

In humans and animals, surprise is a physiological reaction to an unexpected event, but how surprise can be linked to plausible models of neuronal activity is an open problem. We propose a self-supervised spiking neural network model where a surprise signal is extracted from an increase in neural activity after an imbalance of excitation and inhibition. The surprise signal modulates synaptic plasticity via a three-factor learning rule which increases plasticity at moments of surprise. The surprise signal remains small when transitions between sensory events follow a previously learned rule but increases immediately after rule switching. In our model, previously learned rules are protected against overwriting when learning a new rule. Our surprise-modulated spiking network model makes a step towards solving the stability-plasticity dilemma in neuroscience and the problem of continual learning in machine learning.

## Introduction

An event is surprising if it does not match our expectations^1,2,3,4^. The unexpected punchline of a joke^3^, the unexpected continuation of a sequence of tones^5^, harmonies^6,7^ or images^8,9,10^, as well as rule switching such as shift of escape platform in the Morris watermaze^11^ or meaning of cues^12,13,14,15^ induce measurable physiological and behavioral reactions in humans and animals. Without expectations arising from previous experiences, an event such as the observation of a new image may be perceived as ‘novel’ but cannot be ‘surprising’^16,17^.

Surprise is a well-studied phenomenon in the neurosciences^1,2^ and has also been formally analyzed in the mathematical literature^4^. In the neurosciences, startle responses^18^, delayed responses^2^ and pupil dilation^19,20^ are measurable physiological manifestations in response to surprising events. Moreover, EEG, fMRI, MEG and electrophysiological studies show an increase of brain activity shortly after a surprising event^1,21,9,22,23,24,25^. Apart from its potential role for intrinsic motivation^26^, surprise plays a crucial role in learning: surprising events are more memorable^2,27,28^ and allow quick adaptation to a changing environment^29,30^. In this modeling paper, we study the role of surprise in building expectations, modulating learning, and detecting rule switches. Specifically we focus on two aspects. First, surprising events significantly increase the speed of learning^31,32,33,16^ presumably by increasing synaptic plasticity. Second, surprise is involved in the creation and consolidation of memories^2,34,35^, presumably including the memory of rules.

In contrast to mathematical studies that start from a normative framework of surprise^36,37,38,39,40,41,42,43,44,45,46,4^, we take a constructive approach based on a network of spiking model neurons with plastic connections. We consider two aspects of spiking neural networks as crucial requirements for biological plausibility. First, all information about expected and observed events, and an occasional mismatch between the two, needs to be *communicated via spikes*; thus a comparison of subthreshold membrane potentials across different neurons – as required in some existing models^47,48,49^ – is not possible. Second, synaptic plasticity rules should be expressed as *NeoHebbian three-factor learning rules*^50,51,52,53,54,55^ where the changes of a synapse from neuron A to neuron B can only depend on the spikes of neuron A and the state of neuron B (the two ‘local’ factors’) plus one (or several) neuromodulators that play the role of a global feedback signal (third factor) broadcasted to large groups of neurons; in our approach, a detailed synapse-specific feedback as used in the BackProp algorithm^56^ and variants thereof^57,58,59,60^ is not needed.

Our main hypothesis is that surprise manifests itself in a spiking neural network as a *mismatch between excitation and inhibition* in a layer of hidden neurons that represent the current observation and compare it to the expectation arising from earlier observations. Our approach is intimately linked to both the theory of excitation-inhibition balance (E-I balance)^61,62,63^ and the theory of predictive coding^64,65,66,67^.

Predictive coding is an influential theory in the fields of neurosciences^24,64,68,69,70^ and bio-inspired artificial neural networks^71,72,73,74^. In contrast to the classic framework of predictive coding that emphasizes sparsity of activity as a means to minimize redundancy of codes^75^, we emphasize the advantage of predictive codes for generating a surprise signal in spiking neural networks. Importantly, we propose in this paper that *an intrinsic spike-based surprise signal can modulate biologically plausible synaptic plasticity rules so as to achieve fast adaptation and continual learning across rule switches*.

We focus on two related tasks both involving sequences of observations. The first task illustrates the well-known problem of re-adaptation to abrupt switches in the stimulus statistics where the same rule of stimulus generation is unlikely to occur twice^31,42,46^; the second one exemplifies the problem of continual learning across rule switches where each rule should be memorized since it is likely to re-appear^11,15,76^. In both tasks, expectations (’predictions’) must be built by self-supervised learning and change points (’rule switches’) must be inferred from the observation sequence since they are not indicated by a cue. Our model links observations in the neurosciences at the level of single neurons or circuits to psychological phenomena of surprise and provides an alternative to algorithmic approaches to the stability-plasticity dilemma^77,78^, continual learning^13,79,80,76^, context-dependent prediction^81,82,83^, or context buffers in artificial neural networks^72^.

## Results

### Building expectations in a sequence task with rule switching

Imagine the following sequence of numbers

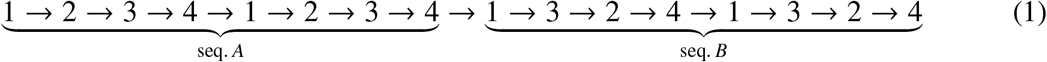

The complete sequence is composed of **transitions** (e.g. 2 → 3) and **switches** (seq. *A* → seq. *B*) between two elementary sequences. The “volatile sequence tasks” used in this paper generalize switches between elementary deterministic sequences, as in (1), to more complex probabilistic sequences generated by a hidden rule (Fig. 1 **A**). We imagine that an elementary sequence arises from a video taken in an empty apartment of ℛ square rooms, each room recognizable by a specific wallpaper. The video camera has been moved around, from one room stochastically to one of the *K* neighboring rooms. In total, ℛ*x*ℛ transitions would be possible, but because of the specific layout of the apartment, not all of these are observed. Note that in a two-dimensional apartment the number of neighboring rooms is *K* = 4, but we sometimes also consider *K* = 2 or *K* = 8 (a determinstic transition corresponds to *K* = 1). While watching the video sequence, the observer sees transitions from wallpapers (’rooms’) 1 → 2 with probability 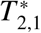 or 1 → 3 with probability 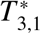, etc. The hidden ‘rule’ of sequence generation arises from the matrix 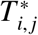that summarizes the layout of the apartment (Fig. 1 **A** and **B**).

**Figure 1:**
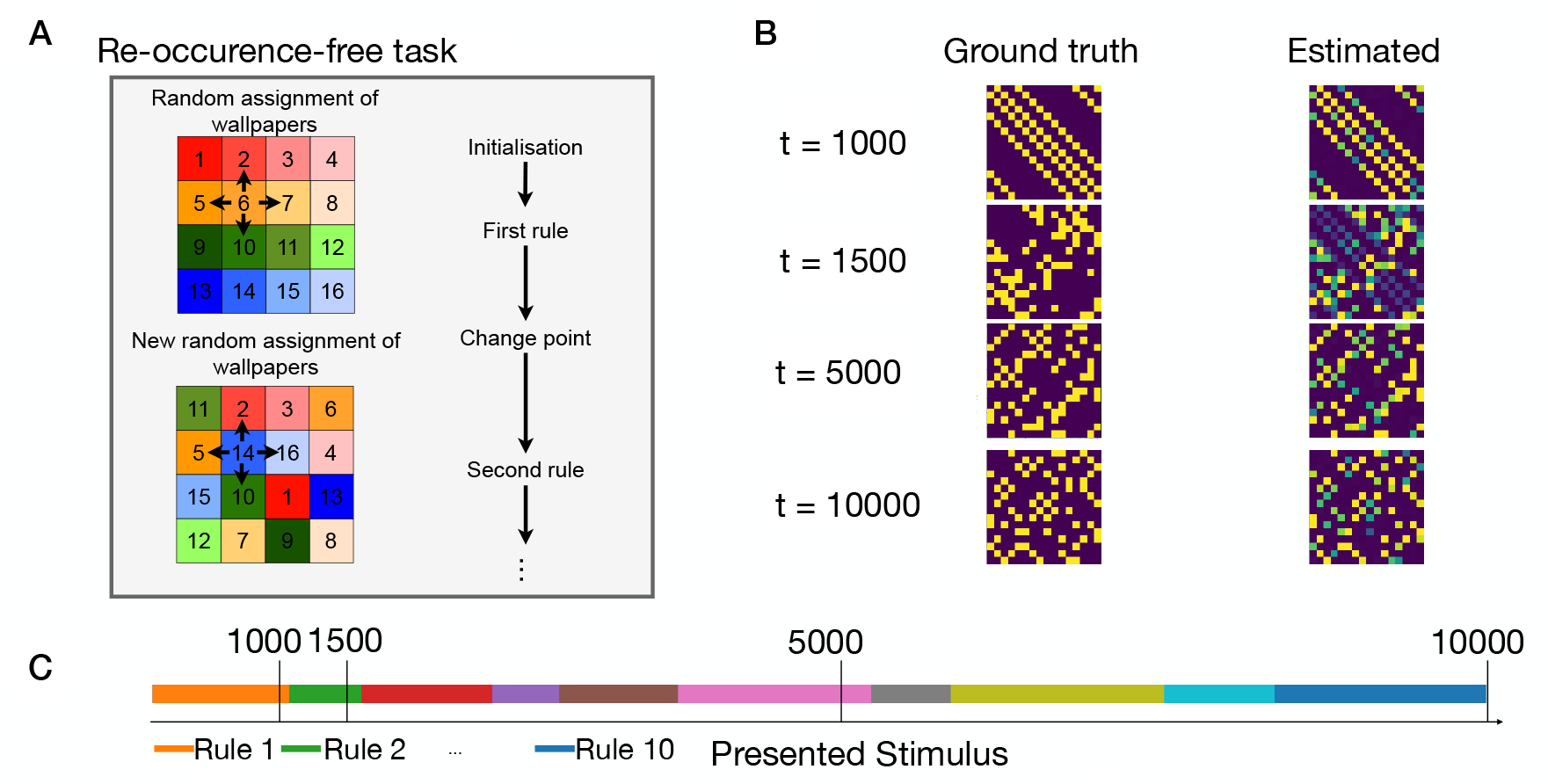
Expected transitions in a volatile sequence task. **A**. At each presentation step of 100ms, the stimulation is caused by the wallpaper (indicated by different colors) in one of the rooms of an apartment with rooms (here = 16). The stimulation sequence reflects transitions (arrows) from the current room to one of the *K* neighboring rooms (here *K* = 4). At rare occasions (change points), the transition rule is changed by a new random assignment of wallpapers to rooms. The same rule is unlikely to return. **B**. The ground truth transition matrix for different rules (left), compared to the transition matrix estimated by the model (right) at different time points of a simulation run. **C**. Switching of rules over time in the simulation of B. Each apartment (Apt1, Apt2, …) only appears once. Vertical lines indicate the time points in B.

In addition to the probabilistic transitions between neighboring rooms, there is occasionally a probabilistic switch (with switch probability ℋ, called ‘volatility’) between different apartments, akin to the switch from seq. *A* → seq. *B* in (1). The ℛ rooms in each apartment use the same wallpapers, but the layout of each apartment is different (Fig. 1 **A**) giving rise to a different rule of sequence generation after each change point^39,47^. The above probabilistic task with rule switching is a generalization of established tasks in cognitive neuroscience of surprise^31,84,85,86^. For example, the task of reference^86^ corresponds to ℛ=2 and *K* = 2.

As opposed to an agent that selects actions to collect information, our observer is passively watching the video with the transitions from one wallpaper to the next. From these transitions between stimuli, the observer learns which stimulus (or stimuli) to expect given the current one, i.e., estimate transition probabilities *T*_*i, j*_. This passive mode is ideal for a study of surprise because, in the context of neuroscience, it avoids any confounding factors arising from action selection^87^ or reward^88^ and, in the context of reinforcement learning theory, it avoids any complex interaction with models of curiosity, action selection policy or questions of model-based versus model-free reinforcement learning^89,90^ - simply because our observer does not choose actions. Once the set of possible transitions in a given apartment has been learned, this knowledge could, of course, be used in model-based reinforcement learning, but this is not part of the tasks that we consider (see Discussion).

A typical sequence of rule switches is shown in Fig. 1 **C** where different rules correspond to transition matrices 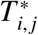 of different apartments. Inspired by experimental observations for passive learning in humans and animals^10,24,85,91,92^, we assume that the (potentially unconscious) goal of observers is to predict possible next observations, i.e., estimate transition probabilities *T*_*i, j*_ that are as close as possible to the real probabilities 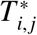. Our spiking neural network model (introduced in the next paragraph) implicitly encodes expectations about possible next stimuli in the set of synaptic weights. From this set of weights we extract the expectations at time *t* in the form of a learned transition matrix *T*_*i, j*_(*t*) that can be compared to the currently active rule 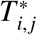 (Fig. 1 **B**). The expectations built during exposure to the sequence are a prerequisite to extract a surprise signal.

### A mismatch of excitation and inhibition yields an intrinsically generated surprise signal

The **Spike**-based **Su**rprise-**M**odulated (SpikeSuM_*rand*_) network model (Fig. 1 **B**) consists of an input layer with random projection onto excitatory and inhibitory neurons in a prediction error layer, and a deep nucleus (i.e., a cluster of neurons in the central nervous system located below cortex^93,68^), e.g., the locus ceruleus^23^, the ventral tegmental area^94^ or higher-order thalamus^95,96^. Neurons in the prediction error layer receive spikes from a first pool of *N* neurons encoding the currently observed stimulus and from another pool of *N* neurons in a memory buffer that encode information on the previously observed stimulus (**Materials and Methods** A.2). Synapses onto excitary and inhibitory neurons have different weights. Two populations of pyramidal neurons *P*_1_ and *P*_2_, putatively located in cortical layers 2/3^24^, compare the weighted inputs of the current observation with the weighted inputs arising via connections from the memory buffer that we interpret as ‘predictions’. Population *P*_1_ is inhibited by the current observation and excited by the prediction coming from the buffer, whereas population *P*_2_ is excited by the current observation and inhibited by the prediction. Both populations project to a group of pyramidal tract (PT) neurons, putatively located in layer 5b^97,55^, which output a low-pass filtered version *Ā* of the summed neuronal activity. Since *Ā* reflects the combined outputs of populations *P*_1_ and *P*_2_, the output of PT neurons can be interpreted as a symmetric measure of ‘distance’ between prediction and observation (**Materials and Methods** A.6). If a prediction is correct, excitation and inhibition balance each other so that the total activity *Ā* of all pyramidal neurons is close to zero.

In our model, the PT-neurons send the filtered network activity information *Ā* to an unspecified nucleus (Fig. 1 **B**) which sends back a neuromodulatory signal 3^*rd*^(*Ā*) that is broadcast across the prediction error layer. We have checked that a large activity *Ā*, caused by positive or negative prediction errors^24,21,98,91,99^ indicates an unexpected transition. A transition is *unexpected* (’surprising’) if the network has for example learned that starting from room ‘10’, the next possible rooms are 1,2,4, or 7 (Fig 1 **A**), but the observed input corresponds to room ‘3’, indicating that a switch point has occurred. Indeed we find that the amplitude 3^*rd*^(*Ā*(*t*)) of the 3rd factor increases after a switch of rules (Fig. 1 **B**, inset). We therefore interpret 3^*rd*^(*Ā*) as a ‘surprise signal’. Note that the surprise signal is a function of activity in the prediction error layer - and therefore implicitly a function of the mismatch between excitation and inhibition.

To achieve E-I balance for *expected* transitions, we assume that activated *excitatory* synapses onto neurons in population *P*_1_ change according to an anti-Hebbian three-factor plasticity rule, modulated by the surprise signal,

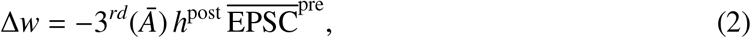

where 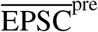 is the filtered sequence of excitatory postsynaptic currents (EPSCs) caused by the presynaptic spike train and *h*^post^ is the input potential of the postsynaptic neuron (for details, see **Materials and Methods** A.4). Analogously, we assume that activated *inhibitory* synapses onto neurons in population *P*_2_ change according to a Hebbian three-factor rule modulated by the surprise signal

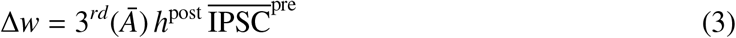

where 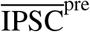 is the filtered sequence of inhibitory postsynaptic currents (IPSCs). For convergence properties of the two learning rules see **Materials and Methods** A.9.

Earlier theories have established that both Hebbian learning of inhibitory synapses^63^ and anti-Hebbian learning of excitatory synapses^100^ lead, for predictable inputs, to a stabilization of the firing rate of postsynaptic neurons at a low value. To check whether this holds also true for the above three-factor rules, we focus on a long stimulation sequence of 3000 presentation steps containing a single switch between two equal-length sequences generated each with a fixed but different rule. Consistent with earlier Hebbian theories, we observe that the SpikeSuM_*rand*_ network converges after about 500 presentation steps to a stationary state of low activity (Fig 2 **C1**). Moreover, the switch between rules causes a sharp peak in the activity *Ā* (Fig 2 **C1**). Thus the activity *Ā* of PT-neurons can indeed be used to extract a surprise signal that is large for *unexpected* observations.

**Figure 2:**
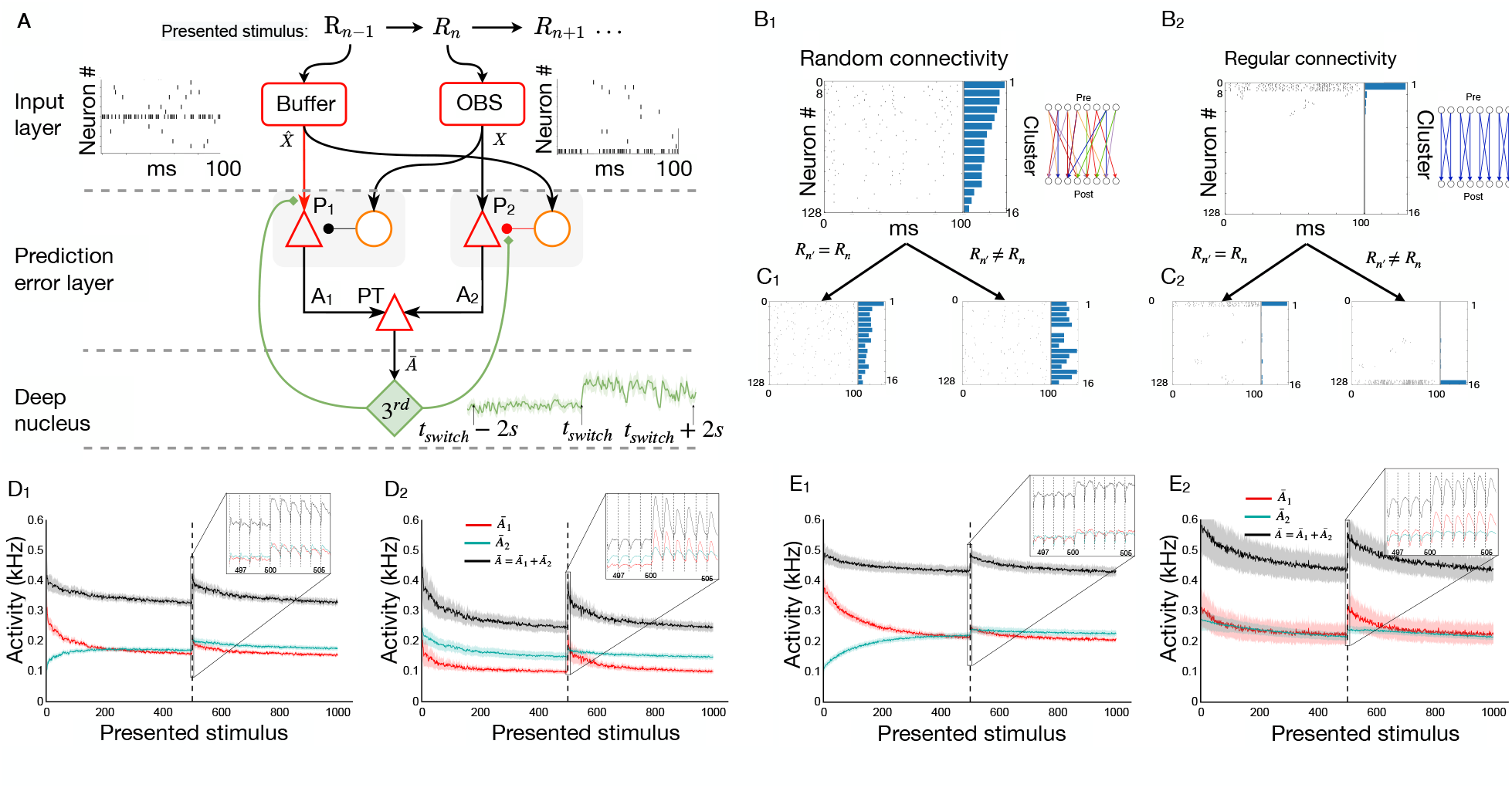
Neurons in prediction error layer exhibit high activity for unexpected transitions. **A**. Spiking network model (SpikeSuM). From top to bottom: Every 100ms stimuli change giving rise to a sequence *R*_*n*−1_, *R*_*n*_, *R*_*n*+1_ … representing wallpapers of the volatile sequence task. The presently observed stimulus (wallpaper of room *R*_*n*_, red box ‘OBS’) and the previous stimulus (*R*_*n* − 1_, ‘Buffer’) are encoded with spike trains of 128 neurons each (16 sample spike trains shown). These spike trains are transmitted to an anti-symmetric excitation-inhibition network (prediction error layer) composed of pyramidal neurons (red triangles) and inhibitory neurons (orange circles). Pyramidal neurons in population *P*_1_ are excited (arrowheads) by the inputs representing the prediction 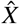 from the previous presentation step and inhibited (round heads) by the current observation *X* whereas neurons in *P*_2_ are inhibited by the prediction 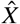 and excited by the current observation *X*. The activity *A*_1_ and *A*_2_ of populations *P*_1_ and *P*_2_ is transmitted to a population of pyramidal tract neurons (PT), which conveys a low-pass filtered global activity 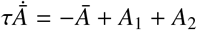 to a group of neuromodulatory neurons in a deep nucleus (green, labeled 3^*rd*^) which send a neuromodulatory surprise signal (inset: Time course of the 3^*rd*^ factor (green) over 4s before and after a switch between apartments at time *t*_*switch*_) back to the prediction error layer. Poorly predicted stimuli increase activity in the prediction error layer and indirectly accelerate, via the 3^*rd*^ factor, learning in the plastic connections (red lines). **B:** Spike trains of all 128 pyramidal neurons in population *P*_2_ during a specific stimulus *R*_*n*_ representing one of the ℛ = 16 wallpapers. For visualization, the 128 neurons have first been ordered from highest to lowest firing rate and then clustered into groups of 8 neurons, with neurons 1 to 8 forming the first cluster, 9 to 16 second etc. The histogram of average firing rate per cluster (horizontal bars) is shown on the right. **B**_1_: Random sparse connectivity from presynaptic neurons in the input layer to neurons in the prediction error layer. Inset: schematics, continuous weights with colors indicating strength from red (weak) to blue (strong); only 8 input neurons (pre) and 8 pyramidal neurons (post) are shown; **B**_2_: Regular connectivity with binary connections. Inset: schematics, nonzero connections (blue) are organized in clusters of 8 neurons, but for readability, only 4 clusters of two neurons each are shown. Random connectivity is potentially closer to biology, but regular connectivity has algorithmic advantages. **C**_1_ **and C**_2_: To compare the response of the network to different stimuli we show the spike train of a new observation *R*_*n*_ while keeping the same order of neurons. For random connectivity spike plots are different if *R*_*n*_ *R*_*n*_ but similar if *R*_*n*_ = *R*_*n*_. The same holds for the regular connectivity scheme, but the structure is more obvious to the human eye. **D**_1_ **and D**_2_: Filtered activity of pyramidal neurons in populations *P*_1_ (red), *P*_2_ (green), and the total filtered activity *Ā* (black) as a function of time averaged over 100 different sequences with a change point (switch of rule) after 500 presentation steps. Both networks indicate a surprising transition (dashed vertical line) by increased activity. After 100 presentation steps, each possible transition (with *K* = 2 allowed next stimuli and *R* = 16 stimuli) has been experienced on average about three times. Insets show the activity before and after the rule switch. **E**_1_ **and E**_2_: Same as in D_1_ and D_2_, but for the case of *K* = 4 possible transitions. Since predictions are less reliable, the activity *Ā* converges to higher levels.

The predictability of the next stimulus is higher in a volatile sequence task with *K* = 2 possible transitions from a given observation than in our reference task with *K* = 4. Hence, the next stimulus becomes ‘more expected’, the prediction error is lower, and the population activity converges to a significatly lower value (Fig 2 **D1**); mean activity averaged over the last 100 presentation steps is 375Hz in Fig. 2 D1 versus 461Hz in Fig. 2 C1, (*p <* 10^−10^). This observation leads to experimentally testable predictions (Supplementary Fig. 8).

We consider two different architectures for the connectivity from the input spike trains to the pools *P*_1_ and *P*_2_ of pyramidal neurons. The first one, SpikeSuM_*rand*_ (Fig. 2 **A1**), uses sparsely connected random projection weights from the input layer to the prediction error layer, whereas the second one (SpikeSuM) has a simplified connectivity matrix with hand-wired binary weights implementing a direct representation of input stimuli by non-overlapping subsets of pyramidal neurons in the prediction error layer (See **Material and Methods A.5**). We find that the qualitative features of the population activity in the simpler network (Fig. 2 **A2-D2**) is similar to that of the randomly connected network (Fig. 2 **A1-D1**). Since results are comparable for the two connectivity patterns, we focus in the following on SpikeSuM with the simple binary connectivity as a reference because it is faster to simulate. Moreover, the interpretation of activity patterns is easier with binary connectivity, since states that are predicted as potential observations given the previous stimulus are identifiable by simple visual inspection of the spiking activity (Supplementary Fig. 9).

### Modulation of plasticity by surprise supports rapid re-adaptation

To understand whether the modulation of plasticity by surprise is necessary for the rapid readaptation after rule switches, we compare the results of SpikeSuM over a long sequence of 10’000 presentation steps with those of two simpler networks with the same architecture but different modulation (Fig. 3**A**). First, we observe that a network with a constant learning rate (no modulation, SNN_*nm*_) either converges after a switch of rules with a short delay (for a fixed high learning rate, an example shown in Fig. 3**A**) towards a high-error solution or converges (for a fixed low learning rate, example not shown) to a low-error solution, but only slowly and with a much longer delay. Second, a network with a simpler modulation SNN_*sm*_ (where the 3^*rd*^ factor depends only on one parameter) shows fairly good convergence but adapts more slowly immediately after a switch. We find that, within the family of tested functions, a 3^*rd*^ factor built of two components as in SpikeSuM (Fig. 3**A**, inset) is necessary to reach adaptation that is both fast and precise, but adding a third component does not further improve learning.

**Figure 3:**
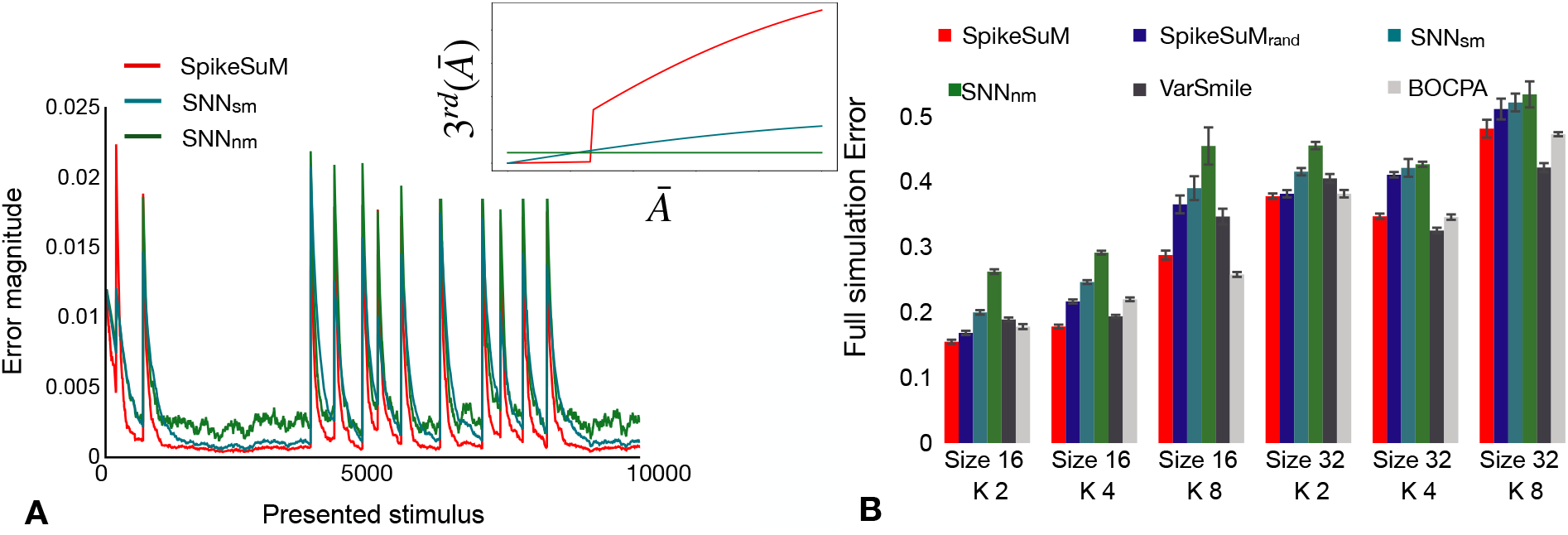
Rapid adaptation enabled by surprise-modulated three-factor plasticity. **A** Inset: The surprise signal transmitted by the 3rd factor as a function of the activity *Ā* in the prediction error layer for three cases (red: SpikeSuM rule; blue: simplified modulation rule; green: constant learning rate, no modulation). Parameters of all three rules have been optimized. Main graph: Error magnitude of the transition matrix (Frobenius norm between the true transition matrix *T* ^∗^ and the estimated matrix *T*) of the SpikeSuM model (red), a Spiking Neural Network model (SNN) with the same architecture and number of neurons, and simple modulation (blue SNN_*sm*_) or without modulation (green SNN_*nm*_), each as a function of time in a volatile sequence task with ℛ = 16 wallpapers and *K* = 4 possible transitions. The occasional abrupt increases in error are due to rule changes in the sequence of observations. The network with surprise-modulated learning leads to faster learning immediately after the switch as well as to better convergence during periods when the rule stays fixed; volatility *H*=0.001. **B** Average error over 10’000 presentation steps with volatility *H* = 0.001 for apartments of different size (16 or 32 rooms) and different numbers *K* of possible transitions from a given room. The performance of SpikeSuM is comparable to that of the Bayesian Online Change Point detection algorithm (BOCPA) and better than SNN_*nm*_ or SNN_*sm*_. The results with random connectivity SpikeSuM_rand_ are below those with regular connectivity, but better than SNN_*nm*_ and SNN_*sm*_.

A systematic comparison shows that SpikeSuM and SpikeSuM_*rand*_ outperform SNN_*sm*_ and SNN_*nm*_ across various instantiations of the volatile sequence task (Fig. 3 **B**). Moreover, the performance of SpikeSuM is only slightly worse than that of the variational Bayesian algorithm varSMiLe^46^ or the online Bayesian change point detection algorithm BOCPA^42^ which are both surprise-based machine learning algorithms designed for near-optimal change-point detection (**Material and Methods** A.13). In summary, on the volatile sequence task without re-occurrence of the same rule, our spiking network with surprise-modulated learning shows faster relearning after a rule switch than the one without which suggests an important role of surprise-modulation in rapid, yet precise, adaptation to changes in the stimulus statistics. Importantly, the surprise signal is not some external variable, but extracted from the spiking activity of the network itself.

### Continual learning across rule switches is supported by the surprise signal

So far each switch of rule introduced a sequence of observations with a new transition matrix so that the previously learned transition matrix could be overwritten at no loss. To explore continual learning across rule switches we now consider a task where the same rules (’apartments’) reappear several times (Fig. 4 **A**). We study a meta-network composed of *M* SpikeSuM modules each acting as one of the rule memories (Fig. 4 **B**). We call this enlarged network SpikeSuM-C (for SpikeSuM with Context). Note that the sets of stimuli (i.e., the different wall papers) are the same in all apartments so that the context needs to be inferred from the observed sequences. Ideally, each module *m* ∈ *M* should focus on one of the context, i.e., a single set of rules. We postulate that in a well-functioning network only predictions within the currently active rule are updated while multiple other contexts that were memorized before are left untouched and can be reused later when the same context reappears.

**Figure 4:**
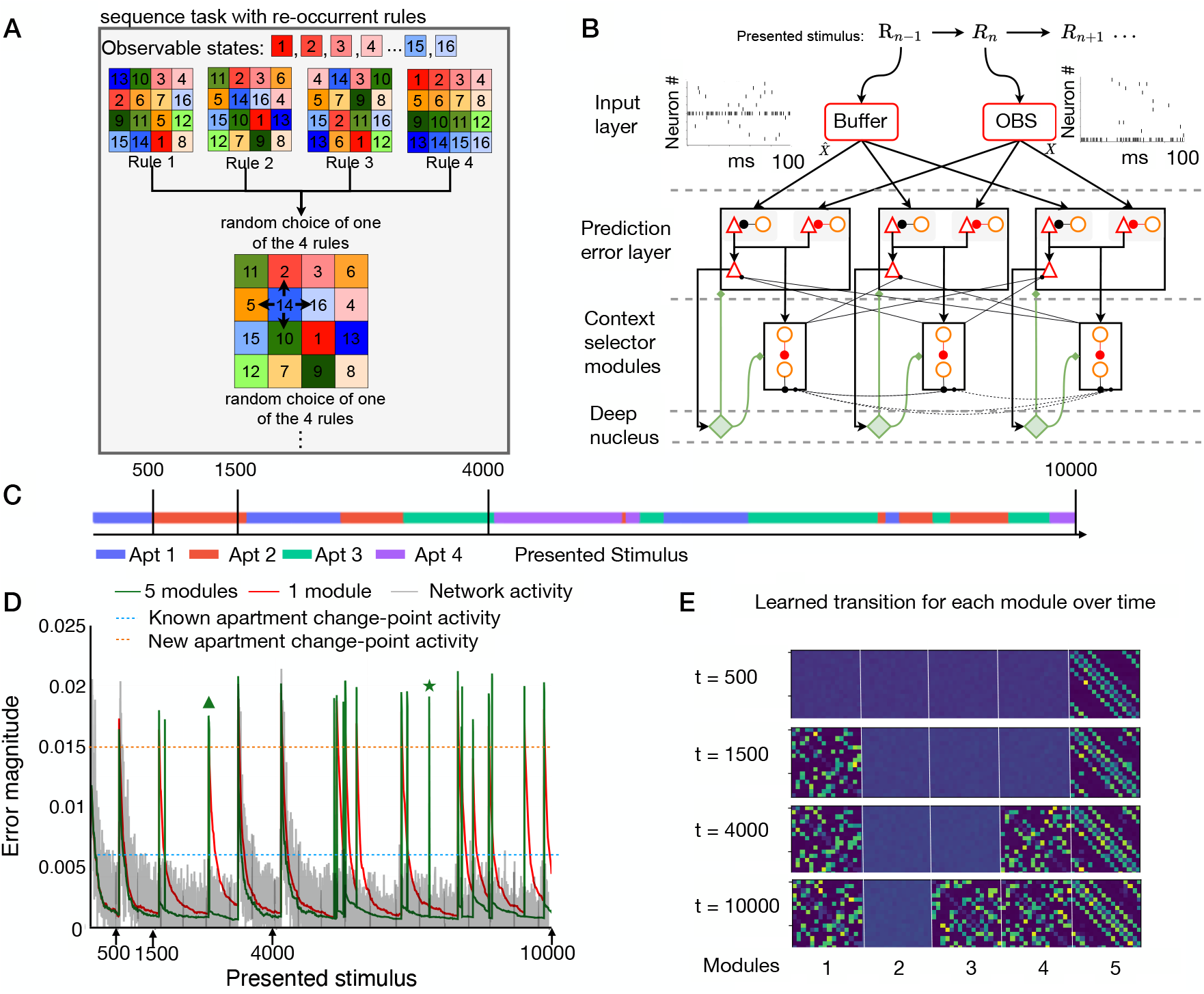
Continual learning across rule switches. **A**: Sequence task with re-occurring rules. Once at the beginning, *M* apartments (here *M* = 4) are created by a random assignment of different wallpapers to ℛ = 16 rooms. At each change point, an apartment is randomly picked from these *M* available apartments, causing a switch of the transition matrix that generates the observation sequence. **B**: The SpikeSuM-C network is composed of four layers. The input layer receives the stimulus and connects to the prediction-error layer which is composed of several SpikeSuM modules (see section); only three are shown. A set of context selector modules (CSM) composed of dis-inhibitory networks is bidirectionally connected with the prediction-error layer. Each SpikeSuM module excites its corresponding CSM. A Winner-Take-All circuit in the CSM layer selects the least excited module. The inhibitory feedback weights from the CSM to the prediction-error layer inhibit the PT neurons of unselected SpikeSuM modules, but not the prediction-error neurons. (see **material and methods A.15**). Red weights are plastic. Non-plastic weights are shown in black for feedforward, solid blue for feedback, and dashed blue for lateral inhibitory connections. **C**: Sequence of rule switches (apartments Apt 1, Apt 2, …) as a function of time. **D**: Summed activity of all PT-cells (grey, arbitrary units) in a SpikeSuM-C network with 5 modules and error magnitude (green, mismatch between transition matrix in currently selected module and ground truth) during learning. When the SpikeSuM-C encounters the second apartment for the second time, the error exhibits a short spike (green triangle) before it instantaneously goes down indicating successful switching between modules. At rare moments (green star marks one of the examples) module switching is initiated at an inappropriate moment but immediately stops thereafter. The activity generated by a rule switch to an unknown apartment is much stronger (grey bars exceed the horizontal orange dashed line) than those to a previously observed apartment (grey bars never reach the cyan dashed line). SpikeSuM-C can create new memories or reactivate memories previously learned if a known transition rule reappears. Red line: behavior of SpikeSuM (control, 1 single module). **E** Evolution of synaptic weight matrices over time for each of the five modules. After 500 time steps the transition matrix of apartment 1 has been learned, and successive transition matrices of other apartments are added as they appear.

To implement this idea, we assume that a set of ‘context selector modules’ (CSMs) selects the specific module that should learn the observed transition (**Materials and Methods A.15**). The indirect coupling of context memories via the CSMs gives rise to a *Best-Predictor-Learns* (BPL) architecture, such that only the context module *m* with the *lowest* activity in the prediction-error layer updates its weights. Importantly, the prediction-error module with the lowest activity is the one with the best prediction for the currently observed transition.

All CSMs compete with each other via standard Winner-Take-All dynamics^101^, such that all CSMs are silent except one. However, none of the prediction error neurons is shut down by the competitive dynamics between CSMs, so that an arbitrary population *p* in module *m* has a non-zero activity. To restrict synaptic plasticity to the prediction error module with the lowest activity, we hypothesize that the nucleus that broadcasts the third factor is organized in several segments, such that segment *m* sends a neuromodulatory signal 3^*rd*^(*Ā*^*m*^) to the corresponding prediction-error module *m*. Such a structure with localized feedback loops is compatible with the anatomy of the higher-order thalamus^95,96,68^ or the ventral tegmental area^94^. More specifically, in our model the activity of populations *P*_1_ and *P*_2_ in the SpikeSuM module *m* excites segment *m* of the nucleus. In parallel, the activity of other CSMs with *m*′ ≠ *m* inhibits segment *m* of the nucleus. Together, excitation and inhibition ensure that only the module *m* with the lowest prediction error updates its weights (**Materials and Methods A.15**)

To illustrate the function of the network, we initialize it with 5 empty context modules and stimulate it with a stochastic sequence generated by switches between four different rules (visualized as apartments Apt1, Apt2, Apt3, Apt4). Fig 4 **D** shows that SpikeSuM-C learns a new apartment as fast as SpikeSuM (equivalent to SpikeSuM-C with 1 module). Moreover, if a known rule reappears it re-activates an existing module instead of learning from scratch. Switches to a previously learned apartment trigger a rapid switch of the network to the correct context. Finally, we find that if the number of learned apartments is smaller than the number of allocated modules, empty modules stay untouched and therefore remain available for later use (Fig. 4 **E**).

The amplitude of the surprise signal after a switch to a previously encountered apartment is smaller than that after a switch to a completely new configuration of rooms (Fig. 4 **D**). In the first case, surprise leads to a switch to an existing module while in the second case to the recruitment of a previously untouched module. Thus, the surprise signals that are self-generated by the network are used by the same network to trigger learning or switching between context modules - all in an unsupervised manner (**material and method** A.16 for more details).

### A modular network architecture avoids the stability-plasticity dilemma

In both SpikeSuM and SpikeSum-C, neurons in the prediction-error layer exibit low activity for known transitions and large transient activities for surprising ones (Figs. 2**C** and 4 **D**). This finding is in agreement with experimental studies using visual^91,102,103^, auditory^85,104,22^, or tactile^105,84^ tasks. The two model networks contain separate populations of neurons, P1 and P2, that respond to negative and positive prediction error, respectively, consistent with in-vivo data^24,21,98,91,99^. Here we ask how plasticity of synapses onto those cells succeeds to avoid the stability-plasticity dilemma^77^ which stipulates that learning is either slow or leads to overwriting of earlier memories.

We find in the volatile sequence task with re-occurrence of rules that the ‘effective synaptic plasticity’ of prediction-error neurons is radically different between SpikeSuM and SpikeSuM-C model networks (Fig. 5). As expected from Hebbian learning, the magnitude of synaptic changes in populations P1 and P2 of the SpikeSuM network *without* modular architecure depends monotonically on the total Hebbian drive, i.e. on the combination of presynaptic activity with an elevated postsynaptic membrane potential (Fig. 5 **A**). Because of the modulation of synaptic plasticity with a third factor signaling surprise, the dependence is even super-linear. In other words, a single event with a large Hebbian drive above a value of 1 (arbitrary unit, a.u.) induces a rapid and strong change of the synapse that causes more plasticity than two events with half the value of Hebbian drive.

**Figure 5:**
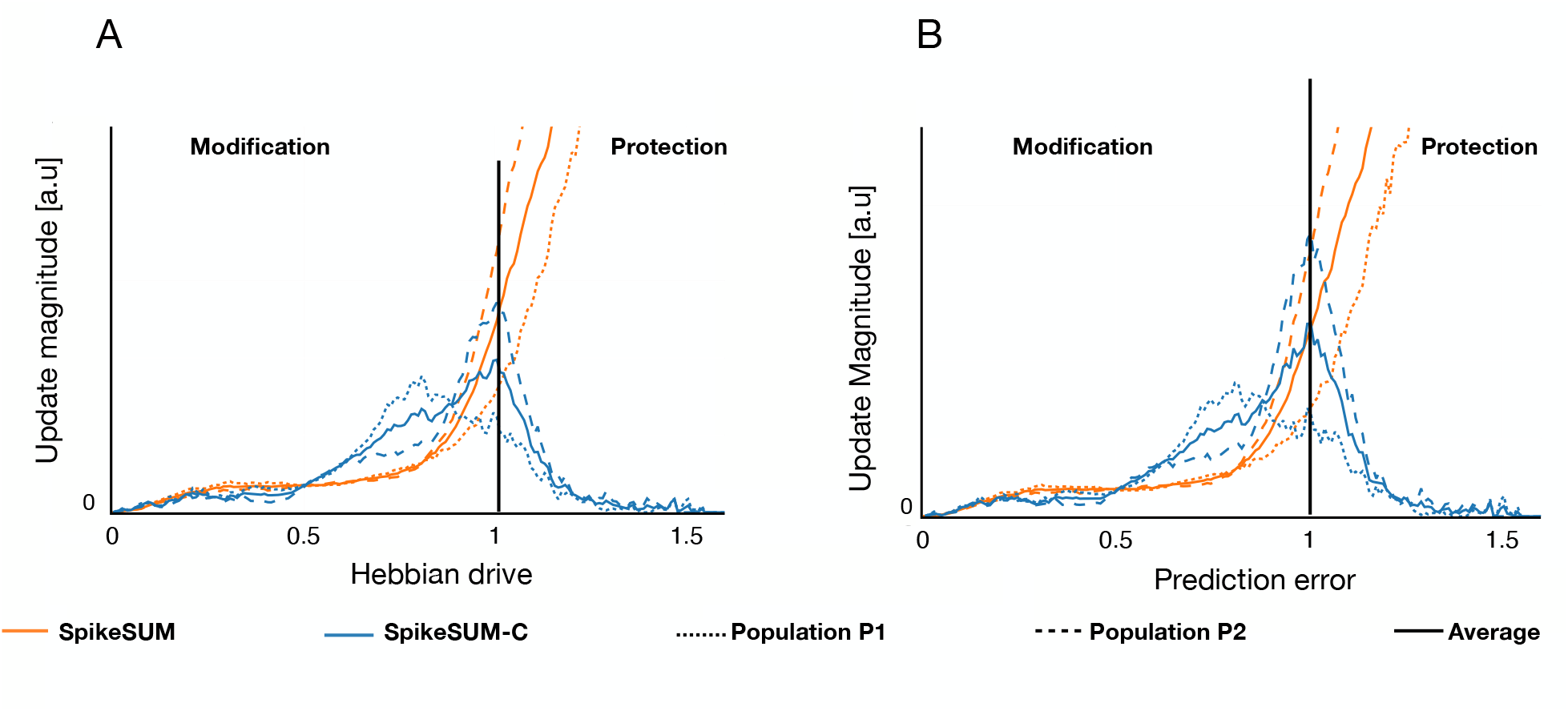
Synaptic Plasticity as a function of Hebbian drive and prediction error. **A**: The magnitude of synaptic change, |Δ*w*_*ik*_| of a specific synapse is shown as a function of the Hebbian drive |*h*_*i*_EPSC_*k*_|, i.e., the combination of postsynaptic membrane potential and the current influx caused by presynaptic spike arrival. The SpikeSuM network (no context detection) is shown in red and the SpikeSum-C in blue. The short-dashed and long-dashed lines show the average over the population P1 and P2, respectively. The solid line is the average over both populations. **B**: The total amount of synaptic plasticity, represented by the update magnitude Σ_*k*_ |Δ*w*_*ik*_| summed over all synapses onto an arbitrary neuron *i* is shown as a function of the prediction error, represented by the momentary membrane potential |*h*_*i*_|, for the SpikeSuM network (red, dashed and solid indicate population averages as in A) and for the SpikeSum-C (blue). For the SpikeSuM network plasticity increases with prediction error and Hebbian drive in both A and B, indicating that large prediction errors systematically induce a large change which eventually leads to overwriting of existing memories after a context change. However, the plasticity in the SpikeSum-C network exhibits two regimes: prediction errors between and 0.5, generate small, but non-negligible changes, and induce a refinement of existing memories, whereas for prediction errors above 1.3 existing memories are protected since other memories are created or changed. The two regimes agree with a prediction of Gershman et al. ^35^.

In contrast, in the SpikeSuM-C network, the dependence is non-monotonic with two different regimes. For small and medium values of synaptic drive, the update magnitude increases with the Hebbian drive whereas, for a Hebbian drive above a value of 1.5 (a.u.), the magnitude decreases back to negligible values (Fig. 5 **A**). The largest update magnitude for a value of the Hebbian drive of 1 marks the transition between regimes of ‘memory modification’ and ‘memory protection’. Memory protection becomes possible because in the SpikeSuM-C network broadcasting of the third factor is not global, but restricted to the currently active network module.

We observe the same qualitative difference between the SpikeSuM and SpikeSuM-C model networks also if we study the total incoming plasticity of a single neuron in the prediction error layer, i.e., the update magnitude of all synapses onto a given neuron as a function of the membrane potential (Fig. 5 **B**). Since the membrane potential encodes positive (in population P2) or negative (in population P1) prediction errors, the graph in Fig. 5B can be interpreted as the total amount of synaptic plasticity (vertical axis) as a function of prediction error (horizontal axis). The small magnitude of synaptic changes for very large prediction errors is functionally important because it leads to the protection of existing modules after a switch of context. The two regimes, i.e. module (*memory*) modification for small prediction errors and module (*memory*) protection for large prediction errors, separated by an inverted-U-shaped curve, have been hypothesized by Gershman et al. as a model for memory modification and formation^35^. Our approach turns a hypothesis at the cognitive level^35^ into specific experimental predictions for synaptic plasticity at the circuit level. In an in-vivo experiment involving multiple contexts, presynaptic activation and postsynaptic membrane potential of putative prediction-error neurons should be monitored while the size of the synaptic connection is measured, e.g., by spine size estimation from optogenetic experiments. We speculate that in primary sensory areas future experimental observations might resemble the qualitative features of SpikeSuM whereas in frontal cortex or subcortical areas those of SpikeSuM-C.

## Discussion

Our network of spiking model neurons enables rapid formation of context-dependent expectations in a paradigm of continual learning where rule switching occur at unknown moments in time. Importantly, rapid adaptation becomes possible by surprise-modulated learning. In contrast to earlier implementations of surprise in cognitive neuroscience models^16,31,32,33,38,40^, surprise manifests itself in our spiking neural model by increased population activity caused by a momentary imbalance of excitation and inhibition^24,63^. The surprise signal has two different roles in our model. First, it triggers the release of feedback signals (e.g., neuromodulators) that serve as ‘third factors’ in an unsupervised neo-Hebbian learning rule^50,51,54^. Second, it initiates switches between modules and avoids overwriting old memories^35,79,80^, since synaptic plasticity is disinhibited only in the module representing the current rule.

In our approach, predictive coding emerges as a necessary byproduct of our aim to extract a surprise signal from spiking activity - as opposed to classic approaches where predictive coding is a consequence of redundancy-reducing or energy-minimizing codes^64,75^. Surprise requires expectations that arise from earlier experience. In our model, the sensory experience of the previous presentation step is represented in the buffer population while predictions are encoded in the connection weights. Our model does not specify whether the buffer population is located in the same area (e.g. cortical L5 cells^24^) or in some other area (e.g., prefrontal cortex^106,107^). The anti-symmetric architecture of the prediction-error circuit in each SpikeSuM-C module requires two separate excitatory and inhibitory pathways that both need to balance each other. This circuit plays a key role in extracting both positive and negative error prediction, similar to putative prediction error neurons in layer 2/3 of the sensory cortex^92,108^. We propose that the activity of these neurons is summed, and potentially low-pass filtered, by layer 5b PT neurons^55,97^ which would then transmit the summary signal (’surprise’) to other areas or nuclei that eventually trigger a feedback signal such as the release of a neuromodulator.

Our model is a conceptual one and makes no specific predictions on the type and/or origin of these feedback signals. However, candidate sources for such feedback signals could be acetylcholinergic neurons in nuclei of the basal forebrain and brain stem potentially linked to arousal and plasticity^68,93,109^, noradrenergic neurons in locus coeruleus linked to cognition, attention, and network reorganization^23,110^, serotonergic neurons in the Raphe nuclei linked to surprise^14^, dopaminergic neurons in the ventral tegmental area related to reward^111^, as well as populations of neurons in the higher-order thalamus potentially related to consciousness or predictive processing^68,95,96^. At least for the last two it is known that the population is not homogeneous but structured^96,94^ which is a necessary condition for the proposed model of switching between different rules encoded in different modules. Even though dopamine is largely correlated with reward and reward prediction error^88,111^, dopamine has also been linked to novelty and potentially surprise^111,112^. The picture is made even more complicated by the fact that dopamine can also be triggered by activity in Locus Ceruleus^113,114^, a nucleus that is traditionally associated with noradrenaline^110^. Hence, a one-to-one mapping between neuromodulators and functional roles should not be expected^52^.

In our model, the surprise signal arises from a mismatch between the representation of the present state and predictions based on the previous state. These learned predictions could be used in several ways. First, the learned predictions generate a synaptic transition matrix that reflects the topology of the input. Topology, which is learned in our model in a completely unsupervised paradigm, forms a compact representation of the hidden structure in the input data that is hard to extract with standard autoencoder models in machine learning^115^. Second, in the absence of input from the next state (e.g., while closing the eyes and waiting), the predictions could be used by an agent to ‘imagine’ the next state - and, if this imagined next state is fed back into the network as surrogate input, it would become possible to plan forward in time over several states, similar to forward replay observed in animals and humans^116,117^ and used in model-based reinforcement learning^89^. Interestingly, since the speed of forward replay would be given by the intrinsic dynamics of the network when closing the loop, it is faster than the original experience^116,117^. Using our model for planning is left for future work.

Predictions in our model are encoded at two levels, i.e., in the weights of synaptic connections and the activity pattern of excitatory neurons in the prediction-error layer (Supplementary Fig. 9). While the model was not designed to reproduce experimental data of frontal cortex neurons, several aspects of the activity patterns in the SpikeSuM-C model are qualitatively consistent with delay activity cortex^107^, implicit encoding of associations^106^, and mixed activity profiles^12,118^ encoding the currently active rule, the present input, the previously active input, and alternative observations consistent with the previous state but inconsistent with the present state.

A distinction between expected and unexpected uncertainty has been proposed in the literature on reward-based learning^33,119^. Analogously, we can define expected and unexpected uncertainty in the absence of rewards. In our volatile sequence task, the expected uncertainty is related to the number *K* of possible next rooms whereas the unexpected uncertainty corresponds to unpredictable switches between apartments. For *K* = 1, the expected uncertainty vanishes. For *K >* 1, the level of expected uncertainty is, after learning, represented in our model by the remaining activity of excitatory neurons in the prediction error layer which could be tested in experiments (Supplementary Fig. 8). Expected uncertainty can also be visible as a non-zero tonic level of the surprise signal (i.e., the 3rd factor). The unexpected uncertainty is represented by sharp peaks in the activity of the prediction error neurons (Fig. 4 **D**).

We view our surprise-detection networks as modules that can be integrated into bigger networks able to solve more complicated tasks. For example, the input layer of our SpikeSuM-C network can be replaced by a multi-layer convolutional neural network previously trained on natural images^120^ with a SpikeSum-C network at the top level, potentially modeling neural activity of the frontal cortex, and other network layers related to areas in the visual processing stream^121^. Moreover, the backprop learning rules of a multi-layer artificial network could be replaced by local predictive learning rules^73^, and SpikeSuM modules can be used in each layer of the network (area of the processing stream) to detect area-specific surprise and locally modulate plasticity. The unsupervised paradigm for training SpikeSuM modules can be complemented by a reward-based paradigm^88^, for example in an actor-critic network^89^. Even though detecting surprise requires a model of possible transitions, recent evidence from human behavior suggests that surprise is mainly used to detect rule changes whereas reward-based learning in complex environments is dominated by model-free reinforcement learning^16^.

Detecting unpredictable switches in the rules governing the momentary environment is a challenge for both artificial neural networks^76^ and biological brains^14,107^. If rule switching is not detected, for example because of reduced serotonergic signaling, behavior exhibits reduced adaptation speed^14^ or even obsessive-compulsive signatures^14,122^. Our model results indicate that surprise signals caused by rule switching manifest themselves as increased neuronal activity and in turn boost learning speed in novel contexts. Surprise in our model is based on prediction errors putatively related to mismatch negativity in EEG signals. Interestingly, schizophrenia patients exhibit a reduced mismatch negativity^123^ and a reduced capacity to make valid prediction^124,125,126^. In our model, missing surprise signals lead to an impairment of memory formation and consolidation, potentially linked to deficits in schizophrenia patients^127,128,129,130^.

Definitions of surprise in a probabilistic framework^4^ have previously been used to explain adaptation to rule switching^42,43,44,45^. However, these definitions cannot be directly applied to spiking neural networks since a correct normalization of probability distributions is difficult to maintain within spiking networks^131,132^ and the calculation of a distance, or Kullback-Leiber divergence, between two probability distributions^4,133^ is even harder. Surprise-driven neural networks for adaptative decision making^134^ or neural particle filters for adaptive perception^135^ are not easily extendable to networks of spiking neurons. Our approach extracts from the activity of spiking neurons a qualitative surprise signal that can be interpreted as a measure of observation-mismatch surprise^4^ without a direct link to probability distributions.

In summary, surprise, i.e., a response of the brain to a stimulus that occurs against expectations^1,2,3,4^, is a phenomenon of a relevance that is comparable to that of reward. Similar to reward and reward expectations^88^, surprise must be detected by neuronal networks in the brain and transformed into modulatory signals that influence synaptic plasticity. Our conceptual model study shows how surprise detection and modulation of plasticity can be implemented in spiking neural networks and how these networks can be used for memory formation, memory protection, and prediction of upcoming inputs, in the absence of supervision or reward.

## Material and methods

### A.1 Two volatile sequence tasks

In the volatile sequence tasks (Fig. 1 **A** and 4 **A**), a sequence of stimuli is generated by a doubly stochastic Markov chain. At each presentation step, a stimulus with index *q* is chosen from a finite set of ℛ different inputs, 1 ≤ *q* ≤ ℛ. Given a stimulus *q* at presentation time step *n*, a stimulus *k* at presentation step *n* + 1 is chosen with probability 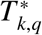 where *T* ^∗^ is the transition matrix that summarizes a given rule. At each presentation step, rules switch stochastically with probability *H* « 1, called the volatility of the rule. We often refer to the moment of rule switch as a ‘change point’. From the point of view of the observer, switches are unexpected and potentially cause a high surprise.

While the theory is more general, we often visualize stimuli as static wallpaper images collected by a video camera that is moved randomly across an apartment composed of ℛ rooms (Fig. 1A), each enabling *K* possible transitions to other rooms. Rooms have distinct wallpapers. The stimulus *R*_*n*_ stands for the wallpaper in the room seen at presentation step *n*. The transitions are stochastic and follow:

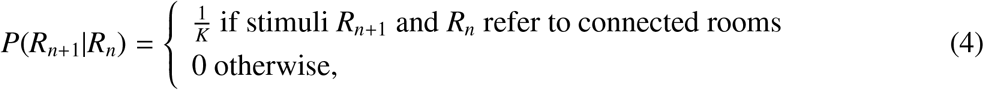

We assume periodic boundary conditions, e.g., room 4 in Figure 1A (top) is a neighbor to rooms 1,3,8, and 16. Thus the layout of the apartment defines the hidden rule of allowed transitions between stimuli. In particular, a transition matrix generated from a given apartment has the property that for each starting stimulus *q*, the elements 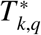 either vanish or take a value 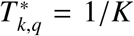 with constraints 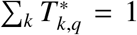 and 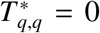. In the theory below we do not assume that the transition matrix is symmetric, even though whenever we simulate an apartment with two-dimensional layout and *K* = 4 (or a 1-dimensional apartment with *K* = 2), then the matrix is symmetric 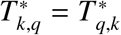.

We design two tasks with different switching patterns. For the first task (’volatile sequence task without re-occurrence of rules’), at each change point, all wallpapers are randomly shuffled. Thus at each change point, a new transition rule is generated. For the second task (’volatile sequence task with re-occurrence of rules’), we first randomly assign the set of ℛ different wallpapers *M* times to different apartments, each of ℛ rooms. At each change point, we randomly choose one of the *M* − 1 possible other apartments. Thus, the number of potential transition rules is finite. The first task implies that having a memory of past apartments is vain as there is a very low probability to observe the same apartment multiple times. Hence, an adaptive algorithm with rapid forgetting is suited to solve this task. For the second task, a suitable algorithm should memorize context dependent predictions and quickly re-activate the correct context after each rule switch.

In the simulations in the main text we use apartments with either ℛ = 16 or ℛ = 32 rooms and vary the number *K* of allowed transitions per room between *K* = 2 and *K* = 8. The terms ‘apartment’, ‘room’, ‘wallpaper’ are for illustration purpose only since each stimulus is represented in the model by a unique neuronal input pattern (see below).

### A.2 Spike trains of sensory neurons

To simulate the volatile sequence task with ℛ discrete stimuli (’wallpapers of rooms’), we translate the wallpaper images (introduced in A.1) into spiking patterns of abstract ‘sensory’ neurons: each stimulus is represented by a distinct cluster of *m* = 8 neurons with elevated firing rate of 100Hz. (We think of these ‘sensory’ neurons as the output of a multi-layer network with wallpaper images as input and 8-hot coding as output; but we do not implement such a preprocessing network). Each stimulus presentation lasts for 100ms (=1 presentation step), and thereafter a new input stimulus is presented to the network. The network input layer is composed of two populations of ‘sensory’ neurons: a population of observation neurons and a population of buffer neurons (see Fig 1). Both populations consist of *m* × ℛ Poisson neurons. Note that we use *m* = 8 neurons per cluster to have a good estimation of the firing rate; however, for a network of rate neurons it would be sufficient to use a single neuron per stimulus (1-hot coding).

In a network of 8 × ℛ presynaptic neurons per sensory population, the first cluster of 8 neurons represents the first stimulus (*q* = 1) of the volatile sequence task, the second cluster consisting of neurons 9 to 16 the second one and so forth. For each observation, neurons in one of the cluster will spike with firing probability 0.1 at each time step of *dt* = 1ms (firing rate 100Hz), whereas all other neurons fire with probability *E* « 0.1 at each time step. Note that, sensory neurons in the buffer population have the same behavior as those in the observation population except that active neurons encode the stimulus number of the previous observation.

### A.3 Transmission from sensory neurons to prediction error neurons

Each spike *z*_*k*_ in neuron *k* of one of the sensory populations triggers a square EPSC of length *l* = 4ms which is transmitted to neurons in the prediction error layer consisting of two populations *p* ∈ {*P*_1_, *P*_2_}. The total input current *I*_*i*_ into neuron *i* of the prediction error layer is

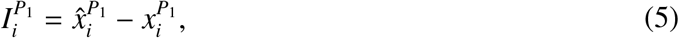

if neuron *i* is in population P_1_ and

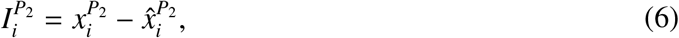

if neuron *i* is in population P_2_. Here,

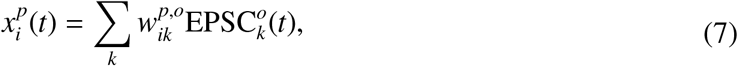

is the input from sensory neurons in the observation population to neuron *i* in population *p* of the prediction error layer where 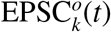 is 1 if neuron *k* in the observation population has fired in the last 4ms and 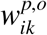 are fixed observation weights. Similarly,

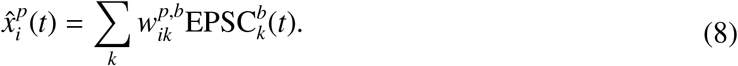

is the input from sensory neurons in the buffer population to neuron *i* in population *p* of the prediction error layer where 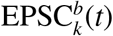 is 1 if neuron *k* in the buffer population has fired in the last 4ms and 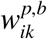 are plastic weights driven by the plasticity rule of Eq. 19. We refer to 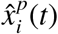 as the (learned) prediction and to 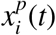 as the (representation of the present) observation. To simplify the notations we drop in the following the time argument *t* and replace

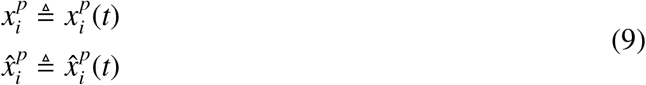

### A.4 Spiking neuron model

Neurons in the prediction error layer are described by the Spike Response Model SRM_0_^136,101^. Each prediction error neuron *i* receives an input current 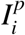 where *p* stands for *P*_1_ or *P*_2_; cf. Eqs. 5 and 6. The input current is then integrated to obtain the input potential (Fig. 6)

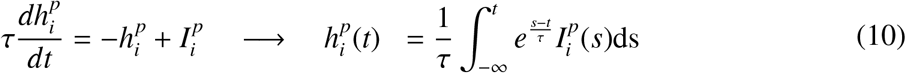

**Figure 6:**
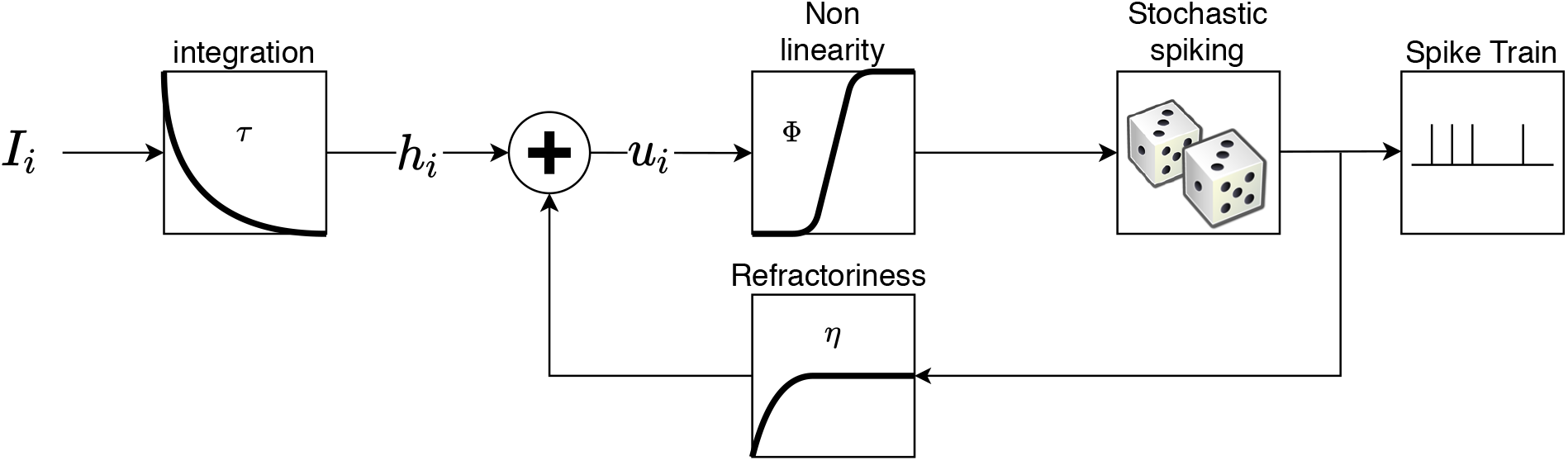
Spike Response Model of neurons in the prediction error layer. Each postsynaptic neuron receives an input current *I*_*i*_. This current is integrated, with membrane time constant *τ*, to obtain the input potential *h*_*i*_. The actual membrane potential of the neuron *u*_*i*_ is the combination of both the input potential and a refractory function *η*, where *η* is a strong negative potential activated after a spike, forcing the neuron to stay silent for a while. The spike times are then randomly drawn with probability *ϕ*(*u*_*i*_) generating the spike train of neuron *i*. Redrawn from ^101^.

Combining the input potential with a refractory kernel *η* leads to the membrane potential

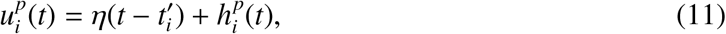

Where 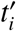 stands for the *last* firing time of post-synaptic neuron *i*, and 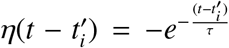 is an exponential refractory function, preventing the neuron to fire again right after a spike. Spikes are generated stochastically with probability

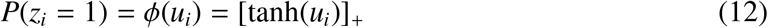

per time steps of *dt* = 1*ms* where *ϕ* is the activation function of the neurons and [*x*]_+_ = *x* for *x >* 0 and zero otherwise. Eqs. (10), (11) and (12) define the Spike Response Model of the prediction error neurons.

### A.5 Two connectivity patterns onto the prediction error layer: random and regular

The first projection pattern (SpikeSuM_*rand*_) is sparse random connectivity (with density 0.1) and weights uniformly drawn between 0 and 1. In other words, for each of the 256 postsynaptic neurons in the prediction error layer we draw an input connection to a specific presynaptic neuron with probability of 10 percent and then connect the two neurons with a random weight (Fig. 2). Since in our standard simulations we have 16 different stimuli and each stimulus is represented by a distinct cluster of *m* =8 presynaptic neurons, the average number of input connections to a neuron in the prediction error layer is 0.1 · *m* · ℛ = 12.8 with a mean weight of 0.5. Thus, in the prediction error layer stimuli (wallpapers) are represented by overlapping groups of neurons of different firing rates (coarse coding).

The second projection pattern is a regularly structured connectivity pattern (SpikeSuM). Presynaptic neurons are, as before, separated in ℛ clusters of *m* = 8 neurons each, but each cluster projects (with binary weights) to a different group of 8 neurons in the prediction error layer. In other words, both pre- and postsynaptic layers are composed of 8ℛ neurons such that different stimuli are represented in the prediction error layer by distinct, non-overlapping groups of neurons (Fig. 2).

### A.6 SpikeSuM Network architecture

Eqs. (5) and (6) show that 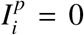 if and only if 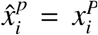, for *p* ∈ {*P*_1_, *P*_2_}. Note that 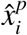 is the prediction arising from the activity of buffer population whereas 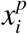 is the present observation. Hence, the total input is minimal if the prediction coincides with the observation. A wrong prediction increases the activity in at least one of the two populations in the prediction error layer: if 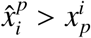 for many neurons in *p* = *P*_1_, then many neurons in population *P*_1_ have a positive input current and nonzero spiking activity; on the other hand, if 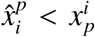 for many neurons in *p* = *P*_2_, then many neurons in population *P*_2_ have a positive input current and nonzero spiking activity. Because of the rectification at the transition from neuronal input 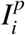 to output spikes (Eq. 12), the two populations *P*_1_ and *P*_2_ complement each other. A natural way to estimate the overall prediction error of the network is therefore to collect the spikes of both populations *P*_1_ and *P*_2_. We assume that the population of PT-neurons acts as a linear filter and transmits a mean activity *Ā* defined as

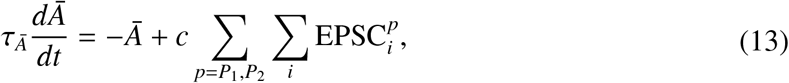

Where 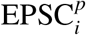 denotes the square excitatory postsynaptic current of neuron *i* of population *p* and *τ*_*Ā*_ and *c* are constants. In our model, *Ā* provides the total drive of neurons in a deep nucleus that receives dense input connections from PT-cells. The neurons in the deep nucleus send back a broadcast signal that measures the total surprise

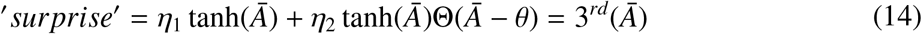

where 3^*rd*^(*Ā*) is a nonlinearly increasing function of *Ā*, Θ is the Heaviside step function and *η*_1_, *η*_2_, *θ* are fixed hyper-parameters. Since this surprise signal modulates learning, we refer to it as a 3^*rd*^ factor that gates plasticity in NeoHebbian three-factor learning rules^54^.

The third factor, composed of two non-linear components, could either be interpreted as a single neuromodulator with a complex nonlinearity, or alternatively as the combined action of two neuromodulators involved in surprise-based learning^33^. Following the terminology of^33^, the adaptation to the *expected* uncertainty (e.g., stochastic transitions to one of the possible next stimuli under a fixed rulen) could be controlled by the action of acetylcholine [described in our model by the term *η*_1_ tanh(*Ā*)], whereas the adaptation speed to the *unexpected* uncertainty (i.e. a rule switch) could be controlled by the action of norepinephrine [turned on in our model if *Ā > θ*].

### A.7 SpikeSuM learning rule: Derivation of Hebbian factors

We aim for a neoHebbian plasticity rule with three factors^54^, i.e., a rule that combines traces of pre- and postsynaptic activity with a modulation of the learning rate. As indicated above, a good prediction of the present observation is indicated by small current in Eqs. (5) and (6) or, similarly, by a small value of the input potential 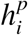 of all pyramidal neurons in populations *P*_1_ and *P*_2_; cf. Eq. (10). We therefore minimize the loss function

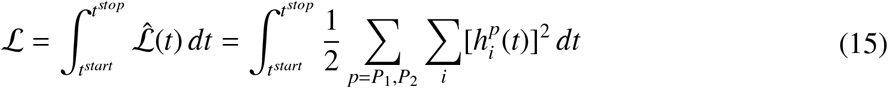

where *t* is time and runs from the beginning *t*^*start*^ to the end *t*^*stop*^ of the experiment. Optimization is implemented as online gradient descent with respect to the weights 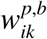 that project from neuron *k* in the buffer population to neuron *i* in population *p* of the prediction error network. We recall that weights 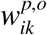 from observation neuron *k* to neuron *i* are fixed. We present here all the calculations for *p* = *P*_1_ only. For the population *P*_2_ one just needs to add a minus sign. The integral over time corresponds to a batch rule; for stochastic gradient descent (online rule) we can focus on an arbitrary point in time and apply the chain rule of differentiation

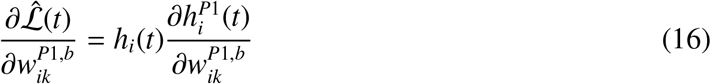

where we can evaluate the derivative using Eqs. (5) and (8)

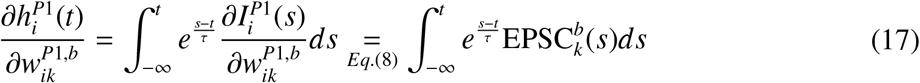

Since EPSCs have a rectangular shape with duration *l* we can evaluate further

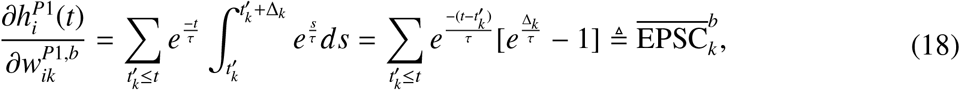

where 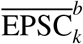 is a low-pass filtered version of the EPSC, 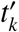 are the spike times of neuron *k* and 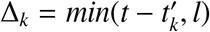. We now apply online gradient descent with an update amplitude proportional to the variable 3^*rd*^ (’learning rate’) and the step size *dt*

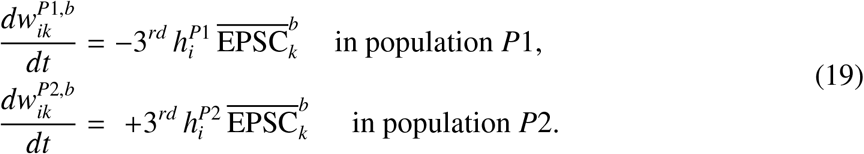

The above neo-Hebbian rule combines a trace of the incoming 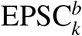 (presynaptic factor) with the momentary input potential 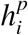 (rather than the spike time) of the postsynaptic neuron (postsynaptic factor): these are the two Hebbian factors. In standard stochastic gradient descent, the learning rate 3^*rd*^ could be fixed or slowly decrease over time as learning proceeds (’freezing’), and also depend (via a momentum term) on the recent history. However, in our model, the learning rate increases whenever the prediction fails (indicated by a large prediction error), so that we refer to the learning rate 3^*rd*^ as a surprise-driven neuromodulator. To summarize, we have a three-factor learning rule with the following properties: (i) EPSC_*k*_ limits the weight update to active connections; (ii) *h*_*i*_ is the local signed prediction error and goes to zero if the prediction for neuron *i* is correct; (iii) finally, 3^*rd*^ is a function of the global unsigned prediction error which is sent back as ‘surprise’ to the full network; see main text and section A.8.

### A.8 SpikeSuM learning rule: third factor

There is no fundamental reason that a learning rate should be fixed as long as each update step (in the batch-rule) decreases the loss^137^. However, in an online gradient descent rule, we have to make sure that all observations get an appropriate statistical weight during the update. In particular, we have to ensure that none of the observations is systematically ignored. This could happen if the learning rate 3^*rd*^ vanished whenever a specific stimulus appears. Such a problem is not a hypothetical one, because of the rectification of the neuronal gain function; cf. Eq. 12. Suppose that for a given stimulus, the observation 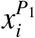 is larger than the prediction 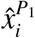 for all neurons in population *P*_1_. In this case, none of the neurons in population *P*_1_ would respond. If we were to use a third factor that is proportional to the activity *A*_1_ of population *P*_1_ (e.g., if we set 3^*rd*^ = *β A*_1_), then this stimulus would never lead to an update.

However, the dependence of the third factor 3^*rd*^(*Ā*) upon the total population activity *Ā* of the prediction error layer together with the anti-symmetric architecture avoids this problem. Whenever the observation does not match the prediction, at least one of the populations, either *P*_1_ or *P*_2_, will be turned on. This is true throughout the simulation because (i) there are many plastic weights that code for each stimulus (e.g., with regular connectivity and ℛ = 16 different stimuli, we have 64 weights coding for each stimulus in each of the two populations); (ii) all synaptic weights in both populations are initialized in the range [0, 1]; (iii) the update rule Eq. (19) is symmetric for both populations (i.e., if the excitatory weights onto a neuron in *P*_1_ increase, then the inhibitory weights onto a neuron in *P*_2_ decrease) which ensures that the symmetries at initialization remain throughout learning.

Thus whenever predictions and observations do not match, the total activity *Ā* conveys a prediction error signal which leads to a non-zero learning rate 3^*rd*^(*Ā*) that is identical for *all* weights.

### A.9 After convergence, synaptic weights of SpikeSuM reflect transition probabilities

We claim that the anti-symmetric architecture of the prediction error layer together with the three-factor learning rule makes the weights converge to a solution that reflects the main features of the hidden rule implemented by transition matrix 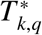 between stimuli. For example, if we imagine an apartment with *K* = 4 possible transitions from a given room, then all four transitions will be encoded in the weight matrix. More generally, we want to show that, for a transition matrix 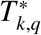 with *K* entries of value 1*/K* per column and zero entries otherwise, the weights onto the pyramidal neurons in the prediction error layer are adjusted such that all possible transitions are predicted proportional to their statistical probabilities. We will also show how to decode the predictions of the network. To stay concrete, we illustrate the abstract mathematical concetps with the words ‘room’ and ‘apartment’, but the calculations are valid for arbitrary stimulus sets and transition matrices.

#### A.9.1 Preliminaries: Encoding of rooms and decoding of predicted transitions

As an abstract encoding of rooms, we use 1-hot encoding. If the total number of rooms is ℛ, then room *q* (with 1 ≤ *q* ≤ ℛ) is encoded by an ℛ-dimensional vector **R**_*q*_ ∈ {0, 1}^ℛ^ with the *q*th component equal to 1 and all other components equal to zero. The transition matrix *T* ^∗^ ∈ ℛ × ℛ that describes the probability of a transitions from room **R**_*q*_ to room **R**_*k*_ has elements 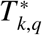 defined as

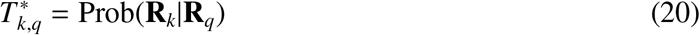

with 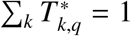 for all *q*. The set of rooms {**R**_1_, …, **R**^ℛ^} represented by 1-hot coding vectors defines an orthonormal basis in an ℛ-dimensional vector space which gives rise to the following properties of the transition matrix *T* ^∗^. First, multiplication of the matrix with the room vectors from both sides gives back the transition

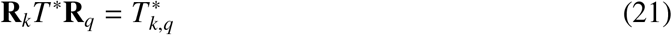

and, second, one-sided multiplication with a room **R**_*q*_ gives a vector **R**_.|*q*_ with non-zero elements for all those rooms that are reachable from **R**_*q*_

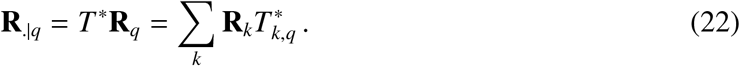

We interpret **R**_.|*q*_ as the code of ‘reachable rooms’ starting from room *q*. It represents the *q*th column of the transition matrix *T* ^∗^ and can be expressed as a linear sum over the one-hot-coded rooms **R**_*k*_. In particular, for *K* = 4 neighboring rooms, the vector on the left-hand-side of Eq. (22) contains four non-zero entries (with a value of 1/4 each) that represent the four rooms reachable from the room with index *q*.

The actual encoding of rooms in the input layer of the SpikeSuM network corresponds to m-hot encoding, since room *R*_*q*_ is represented in the input layer by a cluster of *m* neurons that fire at a high rate (*ν*=100Hz); cf. **material and methods** A.2. For the sake of simplicity of the arguments below, we assume that the neurons representing other rooms *R*_*k*_ *≠ R*_*q*_ are inactive (*ε* → 0) when room *R*_*q*_ is observed. Thus we can think of the input representation of room *q* as an *m*-hot encoding **V**_*q*_ = *P*^1→*m*^**R**_*q*_∈ {0, 1}^*m*ℛ^, where *P*^1→*mℛ*^ is the rectangular expansion matrix from 1-hot encoding to m-hot encoding transforming the the ℛ-dimensional space of rooms into a *mℛ*-dimensional space of input neurons.

We now turn to the representation of stimuli in the prediction error layer. For the SpikeSuM network with *regular* connectivity, the representation in the prediction error layer is also an *m*-hot encoding in each of the two populations *P*_1_ and *P*_2_. However, to keep our arguments general we will also include the case of *random* connectivity. From Eq. (7) we know that the input neurons in the observation population drive neuron *i* in population *p* of the prediction error layer with a current 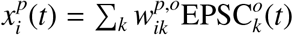. We collect the set of neurons *i* in population *p* into a vector **x**^*p*^, and the weights 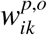 into a matrix *W*^*p,o*^ and write the vector equation

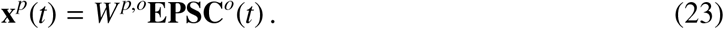

Let us consider a time point *t*_*n*_ located close to the end of the *n*th presentation step. Furthermore let us suppose that during the *n*th presentation step room 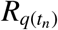 was observed. Here *q*(*t*_*n*_) denotes the index of the room in presentation step *n*.

We exploit the *m*-hot encoding to write for the mean activity pattern in population *p*

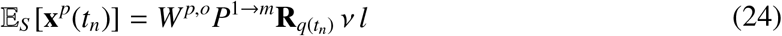

where 𝔼_*S*_ [**x**] denotes the expectation over stochastic spiking of the Poisson neurons in the input layer, *W*^*p,o*^ is the matrix of fixed connectivity weights to the pyramidal neurons in the prediction error layer, *ν* is the firing rate of the active neurons, and *l* is the duration of the rectangular EPSC. Similarly, the expected prediction generated by connections from neurons in the buffer population to those in population *p* ∈ {*P*_1_, *P*_2_} is

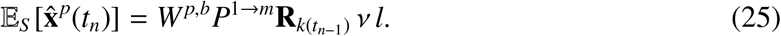

Since we would like to interpret activity patterns in terms of the rooms, we introduce hypothetical decoding weights *D*^*p*^ from the space of neuronal activities (in one of the pyramidal populations in the prediction error layer, *p* ∈ {1, 2}) to the space of room labels in 1-hot coding. We choose decoding weights such that encoding followed by decoding forms an auto-encoder for arbitrary rooms *R*_*q*_:

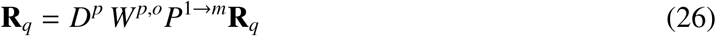

With these decoding weights fixed, the read-out with the matrix *D*^*p*^ enables us to interpret the momentary activity **x**^*p*^(*t*_*n*_) of neurons in the prediction error layer in terms of room labels; to see this compare the right-hand side of Eq. (26) with Eq. (25). Note that the decoding weights are an interpretation tool, but not implemented in the network (even though it would be easy to learn them, for example with the perceptron learning rule).

In order to interpret the *predicted* activity 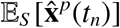 in terms of room labels, we use the *same* decoding weights *D* as for the observed activity

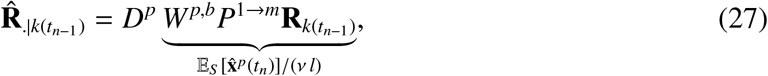

where *k*(*t*_*n*−1_) is the index of the room during presentation step *n* − 1 and 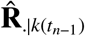 is the prediction of rooms in step *n*, given room with index *k*(*t*_*n*−1_) in step *n* − 1. These predicted room labels enable not only the decoding of predictions in the figures of the results section, but are also at the core of the following theorem.

#### A.9.2 Weights after convergence

We now show the following:

##### Theorem 1.

For a large number of input neurons (*m* → ∞), a small fixed learning rate 3^*rd*^ = *η* « 1, presentation steps longer than the membrane time constant (Δ*T* » *τ*), and a large dwell time in a given apartment (*H* → 0), the synaptic weights connecting the buffer population to the prediction error layer converge under the plasticity rule of Eq. (19) to a fixed point such that (if the input from the momentary observation is blocked) the activity of rectified linear neurons in the prediction error layer can be decoded as

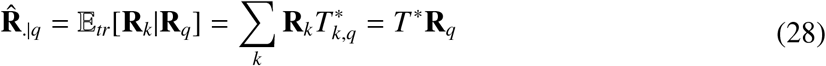

where 𝔼_*tr*_ denotes expectations over transitions conditioned on the index *q* of the previous room. Thus, given that the room in the previous time step *t*_*n*−1_ was **R**_*q*_, the predictive input from the buffer population can be decoded and represents the average of reachable next rooms; cf. Eq. (22)

Notes:

i. For the situation with *K* = 4 transitions per room, the theorem implies that the network activity of the predicion error layer reflects all possible next rooms (with equal weights) if there is no input from the current observation.
ii. The condition of small and constant learning rate ensures a separation of time scales. If learning is slow enough to keep fluctuations of weights small, then learning becomes self-averaging after many presentation steps^138^.
iii. The condition of *m* → ∞ where *m* is the number neurons in the input layer coding for the same room ensures that fluctuations due to spikes, and in particular those correlations between input-and-output spikes that are not accounted for by correlations of firing rates, become negligible^100^.
iv. We only need to calculate the stationary state because for the plasticity rule of Eq. (19) the local stability of the stationary state is guaranteed by^100,63^.

*Proof*. According to Eq. (19) the update of the weights from an input neuron *k* to the set of neurons in population *p* is proportional to the product of the membrane potential and the postsynaptic current PSC(t) (EPSC or IPSC), so that at the end of a single presentation time step of duration Δ*T* » *τ*

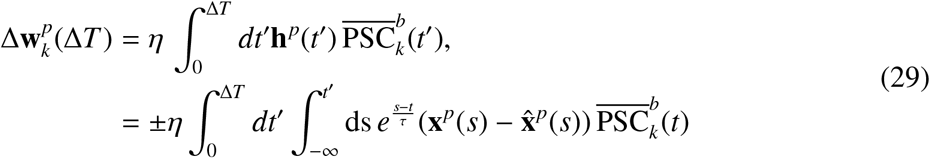

where *η* is a small constant learning rate and 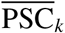 are the filtered PSCs from the presynaptic neuron *k*. The plus-sign applies to population *p* = *P*_1_ and the minus sign to *p* = *P*_2_. We exploited that the presentation time step (Δ*T* = 100 ms) is long compared to the membrane time constant *τ* so that the transients of neuronal activities after the transition between rooms can be neglected.

We study the network at the end of presentation step *n* and assume that during the previous presentation step *n* − 1 the room **R**_*q*_ with index *q* was observed. There are two levels of stochasticity in equation (29), stochasticity of transitions and stochasticity of spike firing. We first take the average over the stochasticity of spiking 𝔼_*S*_, by taking the expectation over the Poisson distribution of input spikes

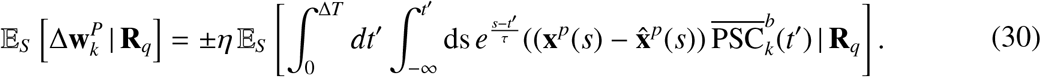

Under the condition *m* → ∞, correlations between input spikes and membrane potential can be neglected^100^. We can therefore separate the conditioned expectations into two independent terms and write

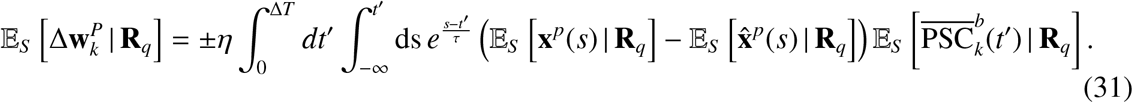

We define the expected input current originating from neuron *k* of the buffer population as 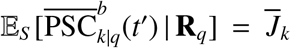 which is constant after an initial transient; this simplifies the notation of the last factor on the righ-hand side of Eq. (31). Furthermore, we use Eqs. (24) and (25) to evaluate the two remaining expectations in Eq. (31).

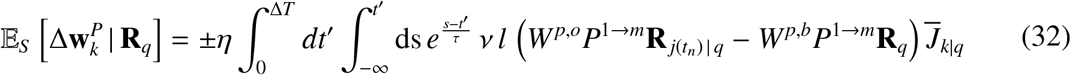

where 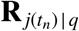 is the room observed in presentation step *n* given that room **R**_*q*_ was observed in step *n* − 1. Note that the index *j*(*t*_*n*_) depends on the specific realization of the stochastic transition starting from room **R**_*q*_.

Exploiting that *H* → 0, we now compute the average over a long observation sequence (expectation 𝔼_*n*_ over presentation steps *t*_*n*_) in the same apartment. We can decompose this average into a multi-plication with the probability *P*(*q*) of observing room **R**_*q*_ and the expected transitions 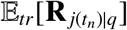 from room **R**_*q*_ to other rooms **R** ^*j*^. We exploit that the rooms reachable from room **R**_*q*_ are given by transition matrix *T* ^∗^.

After convergence, the change of weight 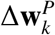 averaged over many presentation steps and realizations of spike trains is zero. Hence we will set 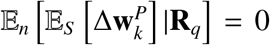. Since the filtering operations induced by the two integrations in Eq. (32) are linear, they yield a fixed factor which can – just like the fixed multiplicative parameters *η ν l* – be dropped after convergence. We exploit that the only term that depends on transitions is **R** ^*j*|*q*^ so that we can pull the transition-average inside and find

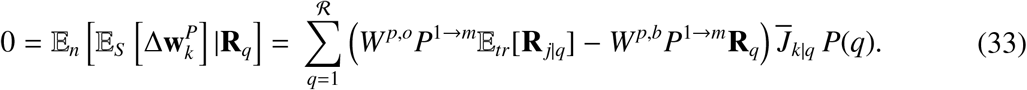

Using Eq. (22) we rewrite 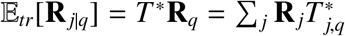.

Since decoding is linear, stationary and deterministic, we can multiple Eq. (33) with the decoding weights *D*^*p*^ from the left. From Eq. (26) and Eq. (27) we obtain

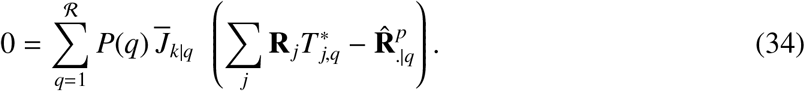

Note that because of the presynaptic factor proportional to 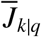, only those weights will be changed that receive input from a neuron *k* coding for room **R**_*q*_. However, since in the long sequence all rooms appear with non-zero probability *P*(*q*) = 1*/K*, for each choice of *k* the synaptic input current 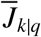 is non-zero during some presentation steps, so that all weights are eventually adapted during the presentation sequence and the terms inside the parenthesis must be zero. Hence

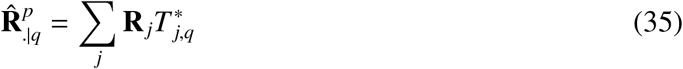

Eq. 35 shows that the synaptic rule with fixed learning rate has a stationary solution where the weight pattern predicts possible next rooms according to the probabilities of the transition matrix. This ends the proof.

Notes:

i. The stationary solution is locally stable both for the plastic excitatory weights in *P*_1_^100^ and for the plastic inhibitory weights in *P*_2_^63^.
ii. In the proof, we decode rooms from the membrane potential of neurons in the prediction error layer. If neurons in the prediction error layer are rectified linear and the input from the observation pool is blocked, then their output is either zero or proportional to their potential. Neurons in at least one of the populations, *P*_1_ or *P*_2_, have a positive potential and can therefore be decoded.
iii. The predictions reported in the Results section are the average across the readouts from two populations *P*_1_ and *P*_2_

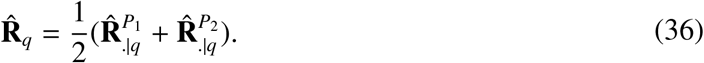

### A.10 Predicted next stimuli with learning rate modulated by surprise

In the previous subsection the learning rate was a constant *η* whereas in our model the learning rate is modulated by the third fact 3^*rd*^(*Ā*). Let us consider a transition from room *q* to one of the reachable rooms. If the reachable rooms have different transition probabilities, e.g, 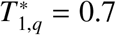 and 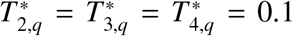, then the transition to room *j* = 1 is less surprising than the transition to one of the other available rooms. Since the amount of activity depends on the surprise level, the third factor will be a function of the room *j* that is reached from room *q*: 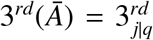. We need to include this dependence in our calculations and modify Eqs. (33) and (34) accordingly. Multiplication on the righ-hand-side of Eq. (33) with 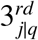 gives a weighted average

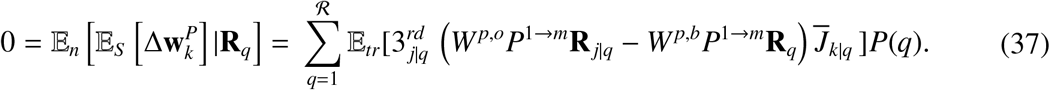

As before, we now use linear decoding weights

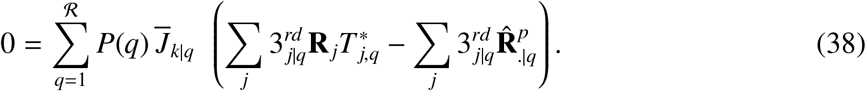

which gives a weighted average for the predicted rooms,

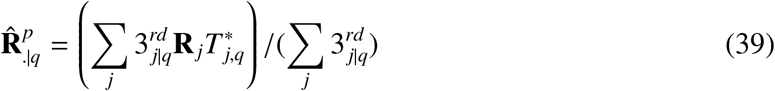

Thus, for surprise-modulated learning rate and inhomogeneous transition probabilities the code of the predicted rooms does not correctly reflect the statistical weights and rare transitions are slightly amplified.

### A.11 Population activity in prediction error layer represents the present state and consistent alternatives to the present state, given the previous state

To illustrate the theoretical results in simulations, we run the volatile sequence task without re-occurrence of rules (Fig 1 **A**) with ℛ = 16 different stimuli (*K* = 4) for 3000 presentation steps in a SpikeSuM network. In the beginning, the spike pattern across the population of pyramidal neurons in the prediction error layer looks noisy (Supplementary Fig. 9, middle left), but after a few hundred presentation steps with a fixed transition rule 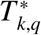 the prediction error layer exhibits four active groups of neurons. These four groups represent the four possible transitions predicted from the *previous* stimulus, including the currently observed one (Supplementary Fig. 9, middle, second panel). Note that the predictions from the memory buffer of the previous stimulus excite neurons in population *P*_1_, whereas the current stimulus mainly excites neurons in population *P*_2_ (cf. Fig. 1). Therefore, with *K* = 4 possible transitions, the currently observed stimulus is represented by a single group of neurons in population *P*_2_ whereas the three alternatives that are consistent with the previous stimulus by groups of neurons in population *P*_1_. Thus, for the SpikeSuM network with hand-wired connectivity, the activity in the prediction error layers reflects a column of transition matrix 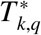where the fixed *q* denotes the previous stimulus (stored in the buffer) and the index *k* runs over the groups of neurons in the prediction error layer coding for stimulus *k*. Immediately after a switch to a new rule, a fifth cluster of active pyramidal neurons is observed. The five clusters correspond to the four wrong predictions (that have been learned with the previous rule now give rise to negative prediction errors) and the currently observed (unexpected) state under the new rule (which gives rise to a ‘positive prediction error’, in the sense that the current sensory input is stronger than the prediction^24^).

### A.12 One-shot learning after a rule switch

As predicted by the theory in the previous paragraphs, the learned weights implicitly encode the full transition matrix of the current rule (Supplementary Fig. 9, bottom). The switch between rules causes a large activity (Fig. 9, top). In turn, the large activity induces an increase of the neuromodulatory surprise signal 3^*rd*^(*Ā*) that leads to a fast update of the weights. Hence the new transition appears in the transition matrix already at the end of the first presentation step, i.e., after only 100ms (red circle in the graph). Thus, a single novel transition is sufficient to change the matrix (learning in ‘one shot’). After spending some time with stimulus presentations under the new rule, the activity *Ā* of PT-neurons returns to a low value and the new transition matrix can be extracted from the weights onto pyramidal neurons in populations *P*_1_ and *P*_2_ (Fig. 9, right, labeled 4).

### A.13 Benchmark algorithms

We compare the results of our network to several state-of-the-art algorithms (Fig. 3). For fairness of comparison, each of these algorithms uses surprise-based online learning. BOCPA^42^ is a Bayesian online algorithm for exact inference of the most recent switch. It is a message-passing algorithm that infers the probability distribution over the run time since the last switch. It is known to be optimal on average for long simulations. VarSMiLe^46^ is a Variational approximation^139^ of BOCPA that uses the Bayes Factor surprise S^*BF* 46^ to detect changes in the environment. VarSMiLe does not need message-passing (as implemented in BOCPA) and has a closed form update rule similar to the SMiLe rule^39^.

We also compare SpikeSuM with simplified networks where we updated the third factor function 3^*rd*^ defined in Eq. (3). The simplification considered are

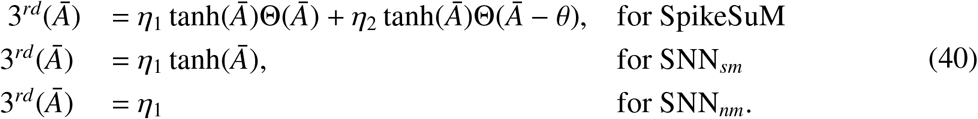

This comparison of 3^*rd*^ factors allows us to investigate the impact of modulation on the learning. The differences in networks parameters can be found in table 1

**Table 1:** Networks parameters for simulations with ℛ stimuli.

| | Presynaptic neurons | Postsynaptic neurons | $3^{rd}$ hyper-parameters |
| --- | --- | --- | --- |
| SpikeSuM | $8 \times \mathcal{R}$ | 128 | 3 |
| SpikeSuM<sub>rand</sub> | $8 \times \mathcal{R}$ | 256 | 3 |
| SNN<sub>sm</sub> | $8 \times \mathcal{R}$ | 128 | 1 |
| SNN<sub>nm</sub> | $8 \times \mathcal{R}$ | 128 | 1 |

### A.14 Simulation parameters and comparison of algorithms

The results are obtained by running networks composed of, 8×ℛ, presynaptic neurons (so that 8 neurons have sustained spiking for each stimulus) and 128 post synaptic neurons (256 for random connectivity). The presynaptic neurons have a firing rate of 100Hz if representing the observed stimulus and the squared EPSCs last for 4ms. The integration time of the input potential *τ* = 10*ms*. See the summary in table 1

Hyper-parameters (*η*_1_, *η*_2_, *θ*) of SpikeSum as well those for SMiLe and BOCPA have been optimized using the python library *scikit-optimize*^140^ minimizing 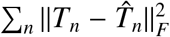, where *T*_*n*_ is the transition matrix of the apartment at time step *t* and 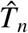 the estimated transition matrix extracted from the weights and ||.||_*F*_ the Frobenius norm.

### A.15 Context Modules: architecture of SpikeSuM-C

SpikeSuM-C is an extension of the original SpikeSuM network and composed of *M* SpikeSuM modules (Fig. 4). Each context selection module (CSM) *m* outputs (in its second layer, L2) an inhibitory current

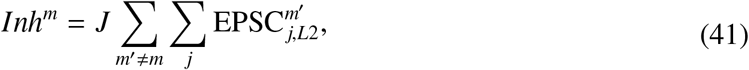

inhibiting the PT neurons,

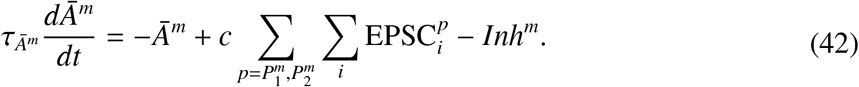

The sum run over all neurons *j* of all CSM other than the one with index *m*. We chose J=20 for strong inhibition. This way if *m* is the active module 3^*rd*^(*Ā*^*m*^) ≥ 0 and 3^*rd*^(*Ā*^*m*′^) = 0.

The neurons in SpikeSuM module *m* are updated following

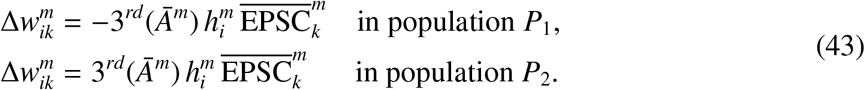

for each SpikeSuM module. 3^*rd*^ is the same as in Eq. (14).

The selection of the learning module is done in the CSM network (Fig. 7) which has a Best-Prediction-Learns (BPL) dynamics. It is a combination of a dis-inhibitory feedforward connectivity within a module and lateral Winner-Take-ALL (WTA) dynamics between modules. Each module contains two layers of *N*^CSM^ spiking neurons each, described by the Spike Response Model in A.4. The first layer (L1) of CSM *m* receives excitatory input from SpikeSuM module *m* and sends inhibitory input to the second layer (L2). L2 is a WTA circuit where the least inhibited module stays selected whereas the other are silenced. The overall network architecture has a BPL-dynamics since the least active SpikeSuM module (i.e., best prediction) is chosen by the CSM network as the learning module. The inputs to neurons in layers L1 and L2 are

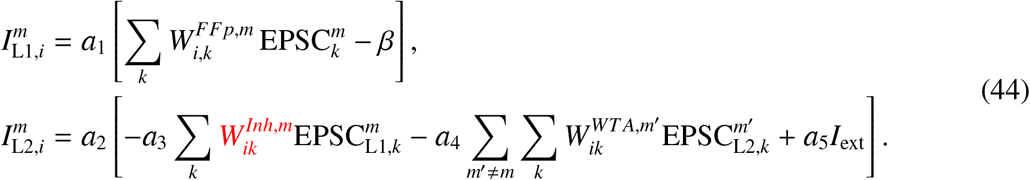

where *a*_1_, *a*_2_, *a*_3_, *a*_4_, *a*_5_ and *β* are fixed positive parameters. Here, *a*_3_ governs the overall strength of the feedforward dis-inhibition drive from the SpikeSuM module. The connection strength *a*_4_ controls the strength of lateral inhibition in the WTA circuit. 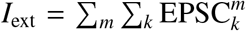, the sum over all possible spikes in the prediction error populations (and all context). 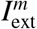 drives the necessary amount of excitation so that the WTA dynamics takes place and accounts for the variability in the network activity, in other words, this noise dependent input allows to stabilize learning, and adapt the WTA threshold. In fact, if the overall activity of the prediction error layer increases due to randomness (not mis-prediction) in the network, the threshold of the WTA would also increase. The parameter *a*_2_ is a scaling parameter that we found useful in setting up the simulations. The choice of parameters is discussed below in **Materials and Methods** A.17.

**Figure 7:**
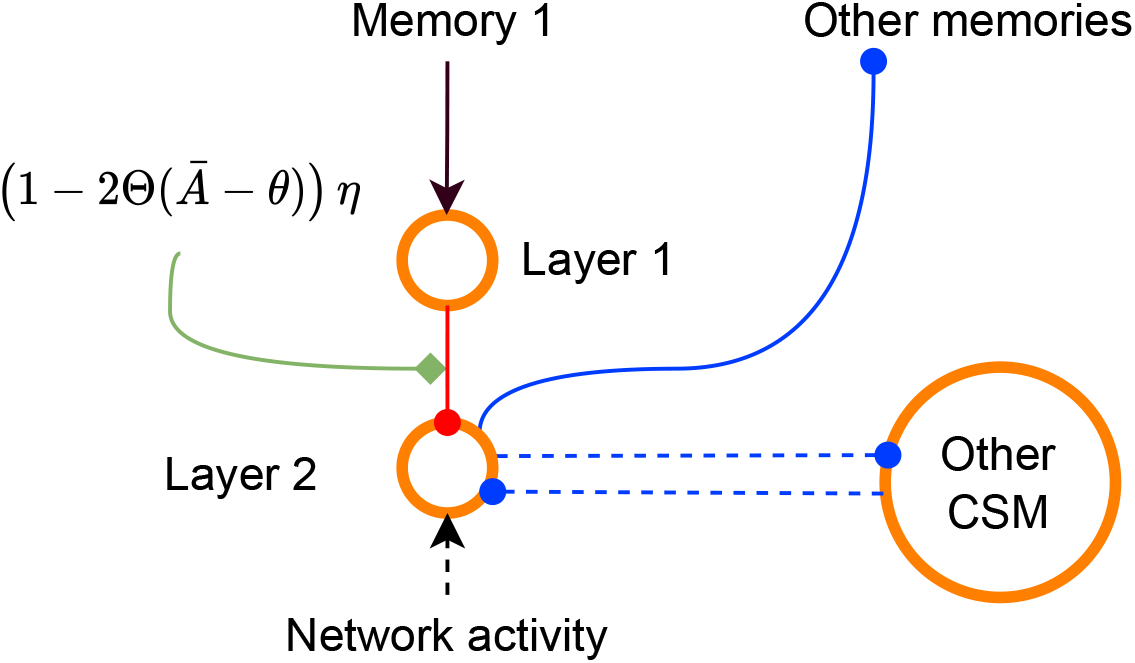
Context selector module (CSM). Each CSM is composed of two layers of inhibitory neurons. Layer 1 receives excitatory input from the corresponding SpikeSuM module. Layer 2 receives inhibition from layer 1 and lateral inhibition from layer 2 of other CSMs. Because of the inhibitory connection from layer 1 to layer 1, the more excitation a CSM receives, the less lower the activity in layer 2. Because of WTA dynamics implemented by lateral inhibition, the CSM module with lowest excitation is selected, inhibits other CSM and shuts down plasticity of other SpikeSuM modules. Red weights are plastic and can be interpreted as ‘commitment’ to the selected module. The network activity is driven by all SpikeSuM modules and supports the WTA dynamic.

While the weights (*W*^*FF*^)^*p,m*^ and *W*^*WT A,m′*^are fixed at a value of one, the connections 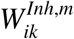 from L1 to L2 are plastic. The inhibitory connections 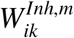 are potentiated by a Hebbian rule modulated by a third factor and depressed by an unspecific decay term with decay rate *α*

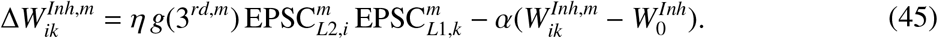

We call these weights the confidence weights. Indeed, 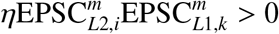 drives the potentiation as long as neurons in both L1 and L2 are active; i.e. when the module *m* is selected. During this phase we consider that the observer builds its confidence up.

After learning a task for some time, 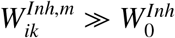 its initial value. Suppose now that suddenly a higher prediction error in SpikeSuM module *m* occurs. This causes an increase of activity in L1 and leads to strong inhibition (because the weights had been potentiated earlier) of neurons in L2 so that the WTA mechanism rapidly ‘un-selects’ this module in favor of another one. Note that postsynaptic neurons in L2 of an CSM that lost the WTA competition are silenced so that connection onto these neurons are no longer potentiated. The net result of the plasticity rule is that modules that have never been chosen in the past have connection weights that are still close to their initial values - and these modules can then be later selected by the WTA dynamics for new tasks. The function *g*(3^*rd,m*^) in Eq. 45 allows to influence the direction of learning. For the simulations we use, *g*(3^*rd,m*^) = (1 − 2Θ(*Ā* − *θ*)) *η* where *θ* is the surprise threshold of SpikeSuM so that weights are depressed during phases of surprise and potentiated otherwise. Finally 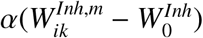, *α* ≪ 1, implements weight decay on a long time scale so that a module that is not used for a long time is slowly forgotten. Note that we could also add a similar decay term to the standard SpikeSuM learning rule in the prediction error layer so as to allow the weights to be slowly forgotten.

### A.16 Systematic results for SpikeSuM-C

To systematically evaluate the performance of the SpikeSuM-C network, we ran a series of simulations for three volatilities H = (0.002,0.001,0.0005). We ran 100 simulations that run over 10 000 presentations, each with 4 randomly drawn apartments and *K* = 2 (or 4) transitions per room. To describe the efficiency of the network we use four different measures of performance:

1. Detection success rate: Number of simulation runs (out of 100) that detect *all* switch points within less than 50 steps and that made less than 10 context switches not linked to apartment switching.
2. Module usage success rate: the match 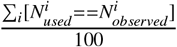 between the number of context modules used in simulation run *i*, 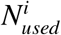, and the number of apartments observed 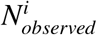.
3. Complete success rate: Number of simulation runs (out of 100) that have detected all switch points in less than 50 steps and that fulfill 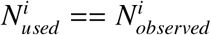.
4. Detection time: mean number of presentation steps between a switch to a different apartment in the input sequence and a switch to a different memory module in the model network.

Note that a simulation where a network uses more memory modules than the objective number apartments does not necessarily imply a complete failure. Indeed, one can conceive that an observer splits a single apartment in two different memories due to the unsupervised nature of the task. This is why we consider in the results section the detection success rate as the most valuable measure (it is not so important how many memories are used as long as one can detect the switch points).

For sequences that switch apartments on average every 2000 time steps 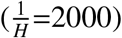 the switch detection success rate is 100 percent for apartments with both *K* = 2 and 4 transitions, but it is slightly lower for simulation paradigms with more frequent context switches (Tables 2a and 2b). We expected that for *K* = 8 transitions the performance breaks down since it implies that from each room and in each apartment 8 out of 16 possible transitions are allowed so that a large fraction of transitions are compatible with several apartments. This was indeed the case (results not shown).

**Table 2:** Summary of SpikeSuM-C accuracy on the re-occurence-based task with different stochasticities and volatilities.

|  | Detection success rate | module usage success rate | Complete success rate | Detection time |
| --- | --- | --- | --- | --- |
| H: 0.002 | $95.0 \pm 2.2$ | $90.0 \pm 3.2$ | $90.0 \pm 3.2$ | $1.4 \pm 0.2$ |
| H: 0.001 | $100.0 \pm 0.0$ | $93.0 \pm 2.6$ | $93.0 \pm 2.6$ | $2.7 \pm 0.2$ |
| H: 0.0005 | $100.0 \pm 0.0$ | $96.0 \pm 2.0$ | $96.0 \pm 2.0$ | $2.3 \pm 0.2$ |

(b) Results for 4 transitions apartment
|  | Detection success rate | module usage success rate | Complete success rate | Detection time |
| --- | --- | --- | --- | --- |
| H: 0.002 | $74.0 \pm 4.4$ | $59.0 \pm 4.9$ | $48.0 \pm 5.0$ | $5.7 \pm 0.7$ |
| H: 0.001 | $98.0 \pm 1.4$ | $91.0 \pm 2.9$ | $91.0 \pm 2.9$ | $6.1 \pm 0.5$ |
| H: 0.0005 | $100.0 \pm 0.0$ | $92.0 \pm 2.72$ | $92.0 \pm 2.72$ | $7.3 \pm 0.83$ |

However, for *K* = 2 already the first or second transition after a switch between apartments is with high probability a good indicator of the switch. Indeed, for *K* = 2 the SpikeSuM-C model detects switches within less than three presentation steps if switches occur on average every 1000 or 2000 time steps (Table 2a).

### A.17 SpikeSuM-C parameters

The results are obtained by running networks with the parameters summarised in tables 3 and 4. The presynaptic neurons have a firing rate of 100Hz if representing the observed room and the squared EPSCs last for 4ms. The integration time of the input potential *τ* = 10ms. The code is available on github (https://github.com/martinbarry59/SpikeSuMNet).

**Table 3:** Summary of SpikeSuM-C parameters for results in table 2a

| Context selection module parameters | 1/H = 500 | 1/H = 1000 | 1/H = 2000 |
| --- | --- | --- | --- |
| $a_1$ | 0.256 | 0.22 | 0.22 |
| $a_2$ | 0.05 | 0.05 | 0.05 |
| $a_3$ | 4 | 4 | 4 |
| $a_4, a_5$ | 20 | 20 | 20 |
| $\alpha$ | 1e-06 | 1e-06 | 1e-06 |
| $\beta$ | 0.045 | 0.045 | 0.045 |
| $\eta$ | 0.0003 | 0.00025 | 0.0003 |
| $W_{max}^{Inh,m}$ | 0.11 | 0.08 | 0.09 |
| SpikeSuM parameters |  |  |  |
| $\eta_1$ | 1e-05 | 1e-05 | 1e-05 |
| $\eta_2$ | 0.005 | 0.005 | 0.005 |
| $\theta$ | 0.45 | 0.45 | 0.45 |
| $W_{init}$ | 60 | 60 | 60 |
| c | 40 | 40 | 40 |
| EI-neurons | 128 | 128 | 128 |

**Table 4:**
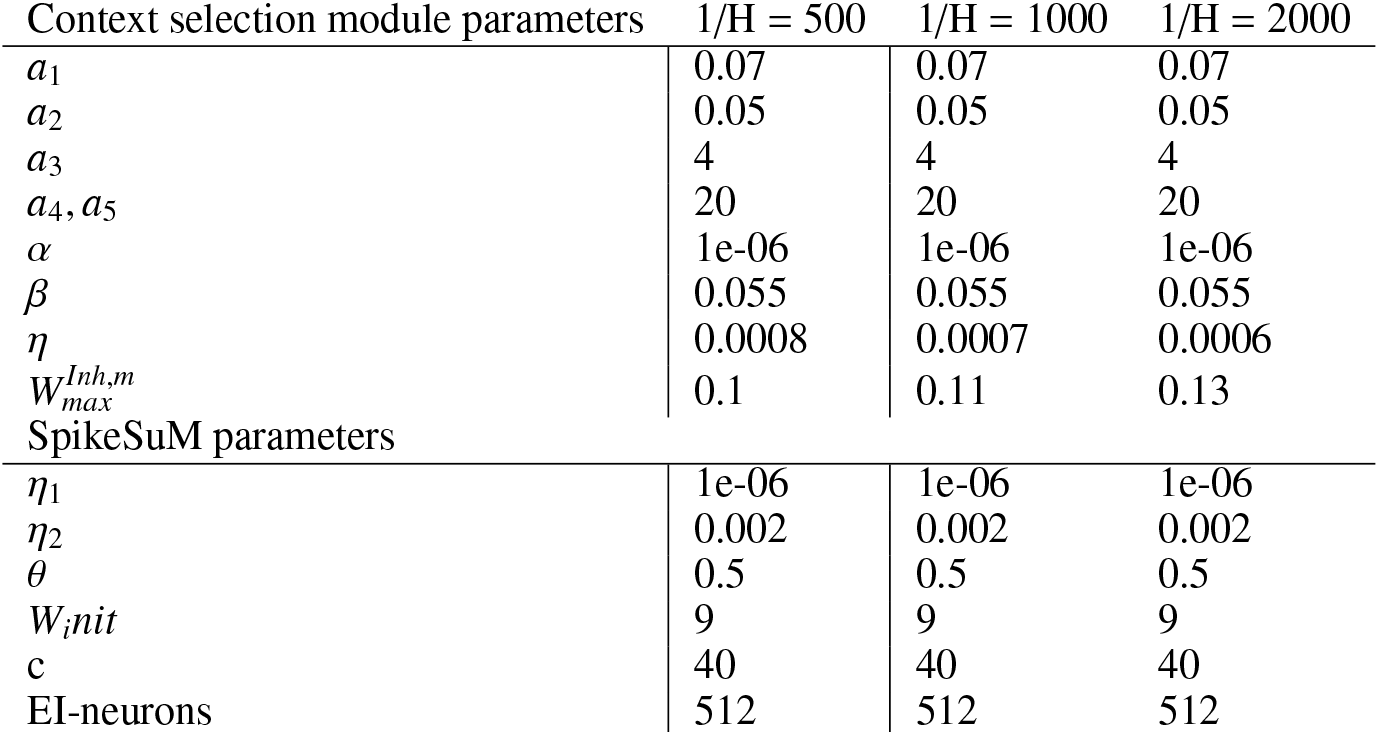
Summary of SpikeSuM-C parameters for results in table 2b

**Figure 8:**
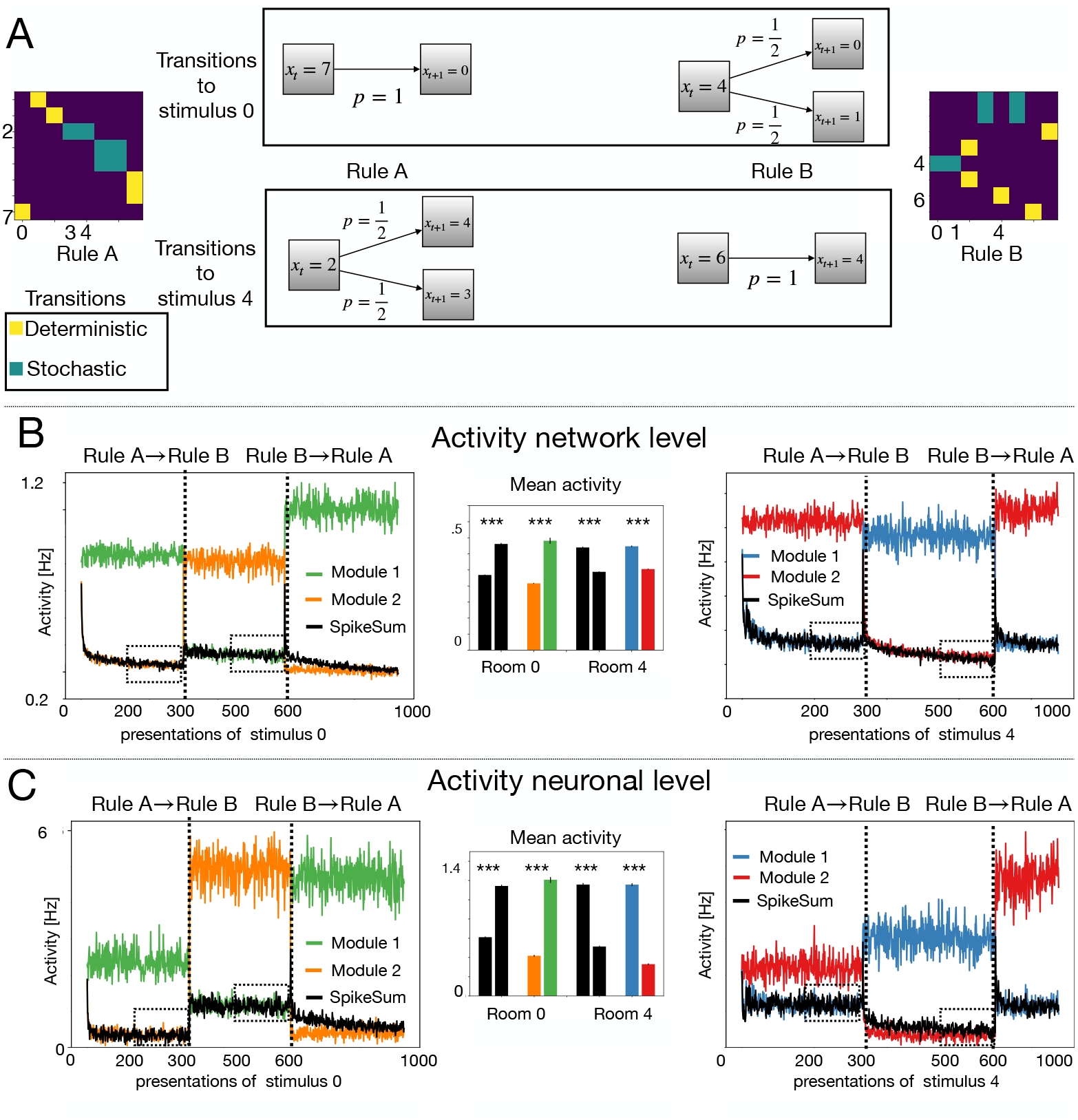
(Supplementary Figure) Deterministic transitions have a different signature than stochastic ones. The paradigm uses a volatile sequence task with re-occurrence of rules but restricted to ℛ= 8 different (auditory or visual) stimuli and two different transitions rules (A and B) and could be tested in rodent experiments. **A** Transition matrix corresponding to rule A (left) and rule B (right). The transition to stimulus ‘0’, *T*_7 → 0_ = 1, is deterministic (yellow square, lower left corner) under rule A but stochastic with a value of *T*_4 → 0_ = 0.5 (light blue) under rule B, and vice versa for the transitions stimulus ‘4’. **B** Population activity averaged over network neurons in populations P1 and P2 during all presentations of stimulus *x*_*t*+1_ = 0 (left) or *x*_*t*+1_ = 4 (right). Black lines: SpikeSuM without context. Green/blue lines: population of neurons in module 1 of SpikeSuM-C (i.e. the module responding to rule A). Orange/red lines: population of neurons in module 2 (i.e., responding to rule B) of SpikeSuM-C. Horizontal axis: count of occurrences of stimulus ‘0’ or ‘4’, respectively. Inset, middle: histogram of average activity after 200 presentation time steps under a given rule. Black bars: comparison of activity under rule A and B in SpikeSuM without context. Colored bars: The activity of neurons in module 1 of SpikeSuM-C during stimuli under rule A is compared with that of neurons in module 2 during stimulie under rule B. A stochastic transition causes more activity than a deterministic one. **C**, same as in **B**, but only the activity of those neurons responsive to stimulus ‘0’ (left) or ‘4’ (right) is shown. In contrast to the simple SpikeSuM network without context, neurons in the SpikeSuM-C network that respond to stimulus ‘4’ in module 1 under rule A (blue) respond even stronger in the context of rule B but this does not affect their plasticity. Thus the same experimental paradigm also differentiate between models with and without context modules

**Figure 9:**
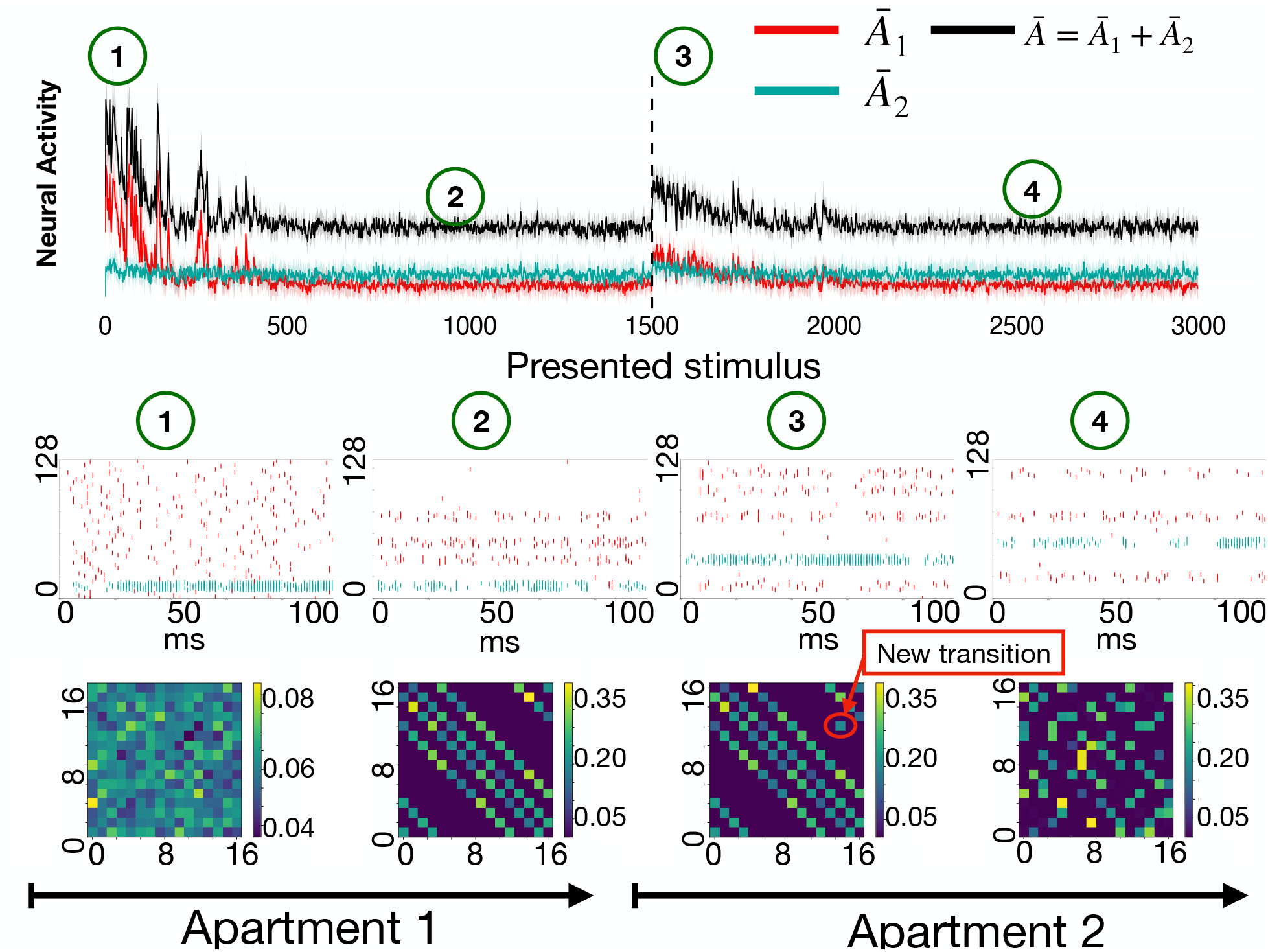
Supplementary Figure. **Mixed response profile where neuronal responses depend on the present state, the previous state, and consistent alternatives to the present state** in a task with ℛ = 16 stimuli and *K* = 4 transitions possibilities and two rules. **Top**: Activity (arbitrary units) of populations *P*_1_ (green) and *P*_2_ (red) as well as the total activity *Ā* (black) of all pyramidal neurons. After 1500 presentation steps, the transition rule switches from rule 1 to rule 2 (Apartments 1 and 2). Each presentation step corresponds to the exposure to one stimulus for 100ms. **Middle**: Spike trains of pyramidal neurons during one presentation step, at different points during learning (from left to right): at the beginning and end of the first episode in Apartment 1 and beginning and end of the first episode in Apartment 2. Initially, (label 1) all pyramidal neurons have approximately the same activity, whereas after 1000 presentation steps (label 2) only a small number of pyramidal neurons are active, i.e., those that correspond to the four possible next rooms reachable from the previously observed room. Out of the four reachable rooms, one is presently observed and, since the observation is stronger than the prediction, neurons in population *P*_2_ fire (blue dots) whereas for the three other predicted rooms the observation is weaker than the prediction so that neurons in population *P*_1_ fire (red dots). After the change in the room configuration (label 3), a fifth cluster of neurons is activated corresponding to the presently observed, but unpredicted, room visible as spikes in population *P*_2_. Pyramidal neurons (16 per room, 8 neurons each from *P*_1_ and *P*_2_) have been sorted according to room number for visual clarity. **Bottom**. Matrix of transitions between rooms decoded from the weights onto pyramidal neurons. At the end of the first presentation step after a change point (label 3), a new element (red arrow) has appeared in the transition matrix corresponding to the newly observed transition, *R*_*n*−1_ → *R*_*n*_. After some time in the novel environment, the new transition matrix is learned (label 4) and the old one is suppressed.

